# Clustering of cortical dynein regulates the mechanics of spindle orientation in human mitotic cells

**DOI:** 10.1101/2023.09.11.557210

**Authors:** Maya I. Anjur-Dietrich, Vicente Gomez Hererra, Reza Farhadifar, Haiyin Wu, Holly Merta, Shirin Bahmanyar, Michael J. Shelley, Daniel J. Needleman

## Abstract

The forces which orient the spindle in human cells remain poorly understood due to a lack of direct mechanical measurements in mammalian systems. We use magnetic tweezers to measure the force on human mitotic spindles. Combining the spindle’s measured resistance to rotation, the speed it rotates after laser ablating astral microtubules, and estimates of the number of ablated microtubules reveals that each microtubule contacting the cell cortex is subject to ∼1 pN of pulling force, suggesting that each is pulled on by an individual dynein motor. We find that the concentration of dynein at the cell cortex and extent of dynein clustering are key determinants of the spindle’s resistance to rotation, with little contribution from cytoplasmic viscosity, which we explain using a biophysically based mathematical model. This work reveals how pulling forces on astral microtubules determine the mechanics of spindle orientation and demonstrates the central role of cortical dynein clustering.

**Highlights:**

- Cytoplasmic viscosity does not determine the spindle’s resistance to rotation
- Each astral microtubule that contacts the cell cortex is pulled on by a single dynein motor
- Pulling forces on astral microtubules determine the mechanics of spindle orientation
- The mechanics of spindle orientation is regulated by clustering of dynein motors at the cell cortex

**Graphical Abstract:** 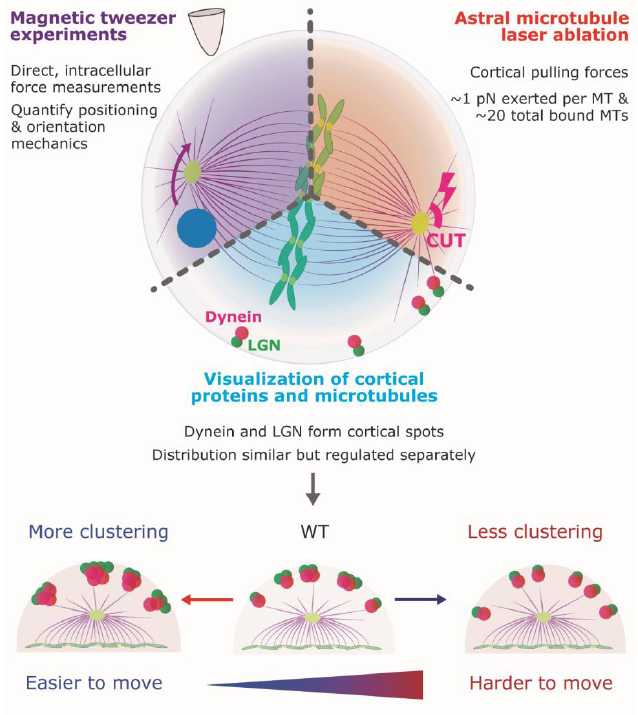

## Introduction

During cell division, the mitotic spindle exerts forces that position chromosomes and segregate them during anaphase ^1^. Forces acting on the spindle dynamically regulate its position and orientation ^2^. The midline of the spindle ultimately sets the division axis of the cell, which is important to building tissue architecture during development ^3^. Thus, the spatiotemporal regulation of forces that position and orient the spindle is crucial to growth and development.

Although much progress has been made in identifying the molecular players involved in successful mitosis ^4–8^, the mechanical processes by which these molecules position and orient the spindle remain unclear ^9^. The most widely accepted model proposes that cortically bound dynein pulls on astral microtubules, thereby exerting forces on the spindle ^10–12^. Previous work has implicated a complex of three molecules involved in recruiting dynein to the cortex—in mammals, membrane-bound gα binds to LGN, which binds to NuMA, which then binds dynein ^12–18^. These molecules appear to form clusters ^8,19–23^, but the precise impact of this clustering on the mechanics of spindle positioning and orientation is unknown.

Beyond the canonical dynein-NuMA-LGN-gα axis, many studies have reported altered spindle orientation when perturbing other cortical proteins, such as DLG1 and MARK2 ^24–28^, or altering microtubule dynamics through TPX-2, GTSE1, and MCAK ^29–31^. Although these perturbations result in increased chromosome and spindle misalignment, the mechanisms by which these proteins contribute directly or indirectly to cortical force generation remains unclear.

There are very few existing measurements of forces exerted on spindles and chromosomes ^32–38^, and none that we are aware of in mammalian cells. The experimental difficulty of direct measurements in live cells, as well as the differences between human and other spindles, have meant that the mechanical aspects of the positioning and orientation mechanisms in human cells remain especially obscure. We have thus far lacked the tools to probe and quantify the mechanical contributions of individual proteins.

Here, we present direct force measurements in human mitotic spindles using a magnetic tweezer system. Combining this with 3D laser ablation and molecular perturbations, we systematically probed the forces that position and orient the mitotic spindle in human tissue culture cells. We characterize the spindle’s motion in response to applied forces, measure its resistance to rotation, and show that cytoplasmic viscosity is not the main regulator of the spindle’s rotational drag. We find that astral microtubules that contact the cell cortex are subject to ∼1 pN pulling forces, consistent with each astral microtubule being pulled on by a single cortically bound dynein motor. Removing dynein from the cortex by *LGN* RNAi greatly reduced the pulling forces on astral microtubules and the spindle’s resistance to rotation, arguing that the spindle’s resistance to rotation is primarily due to astral microtubules being pulled on by dynein. We show that dynein and LGN are clustered at the cortex and that MARK2 and DLG1 regulate both this clustering and the spindle’s resistance to rotation. Finally, we develop a biophysical model of spindle positioning and orientation that incorporates pulling forces on astral microtubules and dynein clustering, which is sufficient to explain the spindle’s resistance to rotation and how it changes in response to both single and double gene knockdowns. Taken together, this work reveals how pulling forces on astral microtubules determine the mechanics of spindle orientation and demonstrates the central role of the clustering of cortical dynein.

## Results

### A calibrated magnetic tweezer system exerts precise intracellular forces on the spindle

To perform direct force measurements on human mitotic spindles, we delivered ∼2.75 μm-diameter superparamagnetic beads to U2OS cells and used home-built magnetic tweezers ^39^ to exert calibrated forces (Fig 1A, S1, S2). Applying a force of ∼15 pN to a spindle pole for ∼1 min caused the spindle to rotate at an approximately constant angular velocity of ω ∼10 deg/min, with the spindle halting its motion and retaining its perturbed orientation after the applied force ceased (Fig 1B,C). Though there was considerable variation in the rotation responses of individual spindles, the trend of a constant rotational velocity during force application (with no rebound) was repeated across cells (Fig 1D).

**Fig 1.**
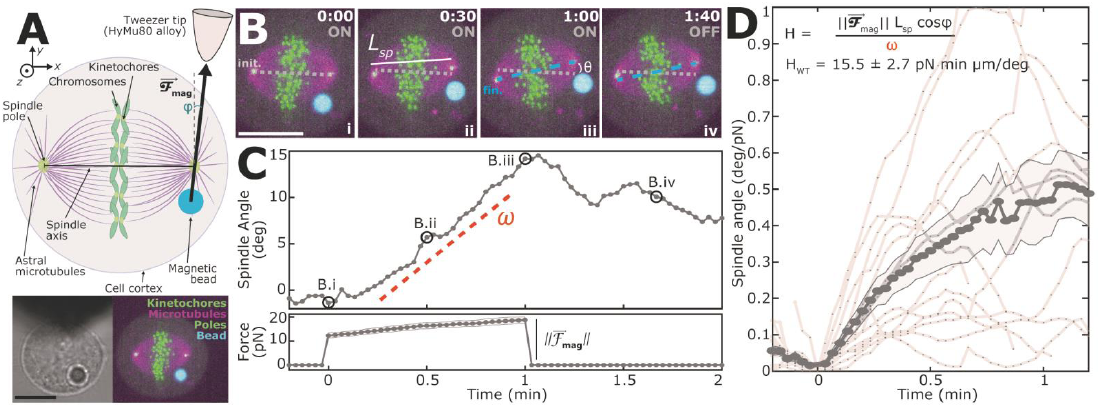
Magnetic tweezers exert calibrated forces directly on the spindle. (A) Experimental setup: an intracellular magnetic bead pushes directly on the mitotic spindle with force magnitude *F_mag_* and angle relative to the spindle φ. Brightfield and fluorescence images of tweezer tip next to U2OS cell (GFP-hCentrin2, green; GFP-CENP-A, green; mCherry-alpha Tubulin, pink) with magnetic bead (647-N biotin, cyan). (All scale bars in this figure are 10 μm. Shaded error bars are SEM.) (B) The spindle rotates in response to applied force. Spindle length (*L_sp_*) and axis angle (θ) marked in panels ii and iii, respectively. Initial (grey) and final (blue) axis angles marked. (C) Spindle axis angle and calibrated force magnitude of B (matching timepoints circled) with angular velocity (ω) and tweezer force magnitude 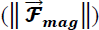 shown. (D) Average spindle rotation response after normalizing to force. The spindle’s resistance to rotation, *H_wt_*, is calculated using ω fit to 0-0.4 min (n=14).

We characterized the spindle’s resistance to rotation by calculating its rotational drag coefficient, *H_wt_* by dividing the magnitude of the torque applied by the bead on the spindle, 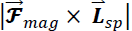, by the average angular velocity, 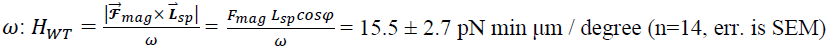 (Fig 1D), where 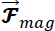 is the force vector applied by the bead, with magnitude *F_mag_* and angle relative to the spindle φ (Fig 1A), and 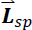 is a vector pointing along the spindle axis with a length *L_sp_*, equal to the average length of the spindle.

### The spindle’s resistance to rotation is not determined by cytoplasmic fluid drag

The simplest hypothesis for the spindle’s resistance to rotation is that viscous forces from the cytoplasm set the rotational drag coefficient ^40,41^. We sought to test this hypothesis in two ways. First, we compared the measured rotational drag coefficient, *H_wt_*, to the rotational drag coefficient predicted based on fluid drag from the cytoplasm alone, *H_cyto_.* We approximated the spindle as a prolate ellipsoid rotating about a pole, giving 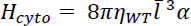, where *l̄* is based on the spindle shape, α is the shape coefficient for a prolate ellipsoid ^42,43^ and *η_wt_* is the cytoplasmic viscosity (*Supplementary Information*). To measure *η_wt_*, we used the magnetic tweezer system to pull a bead through the cytoplasm far from the spindle (Fig 2B) and calculated the displacement and applied force based on the bead’s trajectory (Fig 2C). Using Stokes’s law, we calculated the cytoplasmic viscosity in WT U2OS cells: 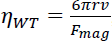, where *r* is the bead radius, υ is the bead velocity, and *F_mag_* is the magnitude of the applied tweezer force, giving a metaphase cytoplasmic viscosity of *η_wt_* = 0.14 ± 0.07 pN min / μm^2^ (n=65) (Fig 2C). The calculated rotational drag coefficient, assuming the spindle’s resistance to rotation comes from viscous force from the cytoplasm, is *H_cyto_* = 3.7 pN ± 1.8 min μm / degree, significantly different from the measured rotational drag coefficient of the spindle, *H_wt_* = 15.5 ± 2.7 pN min μm / degree (p=0.00025), (Fig 2D), indicating that cytoplasmic viscosity is unlikely to account for the spindle’s resistance to rotation.

**Fig 2.**
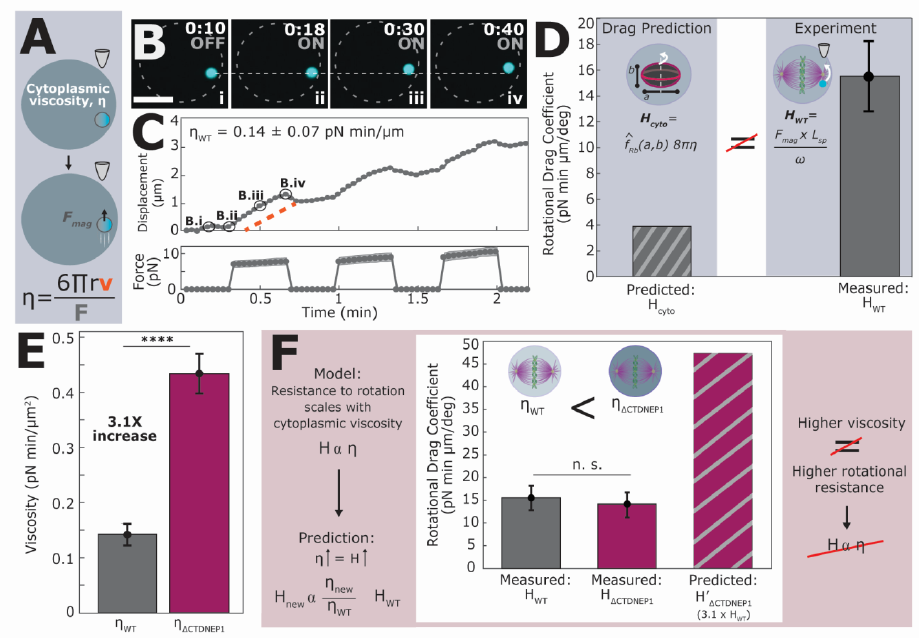
Cytoplasmic viscosity does not determine the spindle’s resistance to rotation. (A) Experimental setup using the magnetic tweezer system and viscosity calculation. (B) A bead (cyan, 647-N biotin) moves in a U2OS metaphase cell (initial position, white). Scale bar is 10 μm. (C) Bead movement and tweezer force from B (matching timepoints circled). (D) The predicted rotational drag coefficient from cytoplasmic fluid drag alone, *H_cyto_*, is lower than the measured rotational drag coefficient, *H_wt_*. (E) The cytoplasmic viscosity of WT U2OS cells (n=65) is increased ∼3-fold in ΔCTDNEP1 U2OS cells (n=39, p<0.0001, ΔCTDNEP1 data also presented in Merta et al., 2021). (F) There is no significant difference between the rotational drag in WT and ΔCTDNEP1 (n=10, p=0.9) cells, and both measurements are lower than the predicted coefficient in ΔCTDNEP1 due to fluid drag.

To further test the contribution of the cytoplasm to the spindle’s rotational drag, we investigated how changing cytoplasmic viscosity impacts the spindle’s rotational drag coefficient. We used a U2OS cell line with a CTDNEP1 knockout resulting in an overproduction of ER membranes ^44^, which had a cytoplasmic viscosity of *η_ΔCTDNEP1_* = 0.43 ± 0.04 pN min / μm^2^ (n=39), 3.1 times higher than WT U2OS cells (p < 0.0001) (Fig 2E, ΔCTDNEP1 data also presented in Merta et al., 2021). If cytoplasmic viscosity were regulating the spindle’s resistance to rotation, then the spindle’s rotational drag coefficient should increase approximately three-fold in the ΔCTDNEP1 cell line (*H’_ΔCTDNEP1_*, Fig 2F, right). Instead, we found no significant difference between the rotational drag in the ΔCTDNEP1 cells (13.9 ± 2.7 pN min μm / degree; n=10) and in WT (p=0.9) (Fig 2F). Taken together, these results show that cytoplasmic drag is not the primary regulator of the spindle’s rotational response.

### Astral microtubules produce net pulling forces on the spindle

Given that fluid forces from the cytoplasm are not the main determinant of the spindle’s rotational drag, another hypothesis is that this resistance is set by interactions between astral microtubules and the cell cortex. We first sought to characterize the nature of the forces that astral microtubules exert on the spindle by cutting them with a laser: if astral microtubules were predominantly pushing, then the spindle would move toward the cut; if astral microtubules were predominantly pulling, then the spindle would move away from the cut. We cut a curved plane with a radius of 2 μm, height of 5 μm, subtending 0-60° off the pole-pole axis with a custom built, reduced repetition rate, femtosecond laser ablation system ^45–47^ (Fig 3A), thereby severing astral microtubules in one quadrant of the spindle, and tracked the subsequent motion of the spindle poles (Fig 3B,C). Spindles rapidly rotated away from the cut astral microtubules (Fig 3D, n=31), while control ablation experiments performed in the cytoplasm near the middle of the spindle (to avoid astral microtubules) produced no significant spindle rotation (Fig 3D, n=13). Therefore, astral microtubules exert net pulling forces on the spindle.

**Fig 3.**
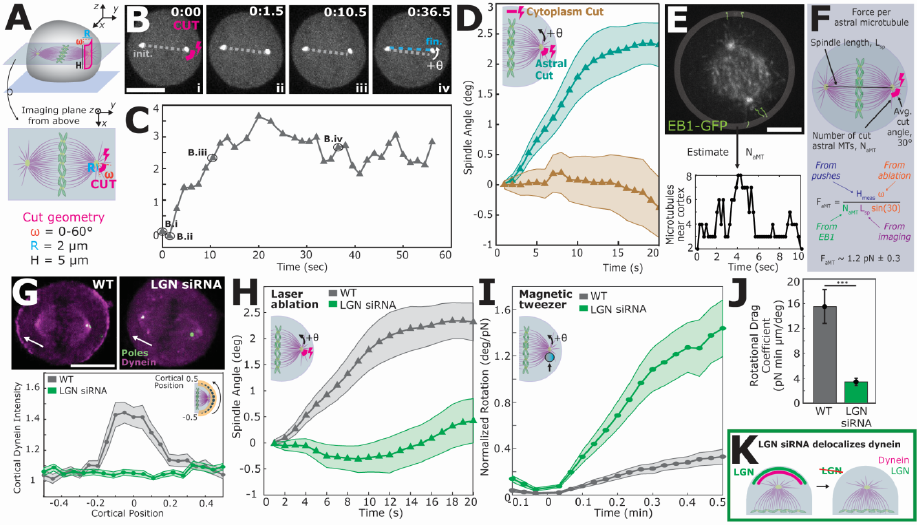
Astral pulling forces strongly contribute to the spindle’s resistance to rotation. (A) Laser ablation removed one quadrant of astral microtubules using a curved plane (2 μm radius, 5 μm height, 0-60° off spindle axis). (B) Spindles rotated after astral microtubules were cut (initial angle in grey, final in blue, cut in pink). All scale bars in this figure are 10 μm. (C) Spindle axis angle after cut (matching timepoints circled). (D) Astral cuts in WT cells (n=31) result in strong rotation away from cut site, compared to no rotation after cuts in the cytoplasm (n=13). (Cut locations and angle sign convention shown in inset) (E) EB1-GFP comets at the midline and cortex (Fig S3E) were used to determine the number of microtubules in contact with the cortex. (F) We calculate the force exerted by each astral microtubule by combining the spindle’s rotational resistance from magnetic tweezer experiments, the spindle’s rotational velocity after astral ablation from laser ablation experiments, and the number of ablated astral microtubules from EB1 measurements. (G) WT U2OS cells treated with *LGN* siRNA had no midline dynein signal (tdTomato-DYNH2), as measured by integrated dynein intensity I_DYN_ (WT 5.26 ± 0.85, n=30; *LGN* 2.26 ± 0.49, n=24, p=0.007). (H) U2OS cells treated with *LGN* siRNA also did not rotate after astral microtubule ablation (n=15), in contrast to WT (n=31). (I) *LGN* RNAi treatment (n=10) resulted in more spindle rotation during magnetic tweezer experiments. (J) *LGN* siRNA treatment significantly reduced the spindle’s rotational drag coefficient by almost a factor of 5 (p=0.0004). (K) *LGN* siRNA treatment removes cortical dynein and reduces cortically-generated force.

### Each bound astral microtubule exerts ∼1 pN pulling force on the spindle

Immediately after laser ablation, spindles rotated away from the cut with an average rotational velocity of *ω_ablated_* = 2.42 ± 0.63 deg/min (n=31). The magnitude of force required to rotate the spindle with that rotational velocity is 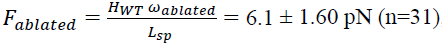, which is presumably exerted by the remaining, non-ablated astral microtubules. Since the spindle does not rotate this way in the absence of ablation, the ablated astral microtubules must normally be exerting an equal and opposite force to prevent such a rotation. Converting this total force to a force per astral microtubule requires knowing the number of astral microtubules contacting the cell cortex in U2OS cells. To this end, we imaged and tracked EB1-GFP comets ^48^ in both the spindle midline (Fig 3E) and at the cell cortex (Fig S3E). For both acquisitions, we calculated the area density of microtubules at the cortex (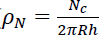, where *N_c_* is the number of microtubules close to the cortex, *R* is the cell radius, and ℎ is the approximate z dimension of the confocal volume) and multiplied that by the total cell surface area, *C_SA_*, = 4π*R*^2^. At the midline, we found an average area density of 0.02 ± 0.008 microtubules/μm^2^, while at the cortex we found 0.04 ± 0.01 microtubules/μm^2^ (n=18). Therefore, we estimate that at any given timepoint, ∼15-45 microtubules are directly contacting the cortex, *N_aMT_*, = *ρ_N_C_SA_*. Assuming approximately 20 astral microtubules and an equal number of microtubules per spindle quadrant, we estimate that roughly *N_aMT,ablated_* = 5 astral microtubules were ablated in our experiments. Combining the number of ablated astral microtubules with the total force exerted by those astral microtubules allows us to calculate the force exerted per astral microtubule on the spindle as 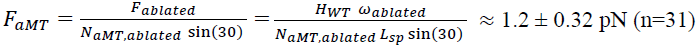 (Fig 3F), where sin(30) accounts for the average angle of the ablated microtubules relative to the spindle axis. This force exerted per astral microtubule is similar to the stall force of human dynein previously measured *in vitro* ^49–54^, suggesting that each astral microtubule that contacts the cell cortex is pulled on by a single motor.

### Dynein is the primary determinant of astral microtubule pulling forces and the spindle’s resistance to rotation

We hypothesized that the pulling forces exerted on astral microtubules were generated by cortically localized dynein. To test this possibility, we investigated the impact of knocking down *LGN* by RNAi, which, consistent with previous results ^12,14,17,18^, significantly reduced dynein at the cortex (with the integrated cortical intensity of dynein, I_DYN_, decreasing from 5.26 ± 0.85 (WT) to 2.26 ± 0.49 (LGN) (p=0.007); Fig 3G) and eliminated the WT phenotype of cortical dynein crescents (Fig 3G, WT n=30, *LGN* n=24). *LGN* RNAi spindles exhibited strongly reduced motion in response to laser ablation of astral microtubules (Fig 3H, WT n=31, *LGN* n=15), suggesting a substantial decrease in pulling forces from astral microtubules. In magnetic tweezer experiments, we observed that *LGN* RNAi spindles rotated substantially more in response to applied force (Fig 3I, WT, n=26; *LGN* siRNA, n=26), resulting in an approximately 5-fold reduction of the rotational drag coefficient to 3.4 ± 0.6 pN min μm / degree (n=9, p=0.0004) (Fig 3J), which is similar to the rotational drag expected solely from cytoplasmic viscosity (*H_cyto_* = 3.7 ± 1.8 pN min μm / degree, p=0.58). These results strongly argue that the pulling forces exerted on astral microtubules result from cortically associated, LGN-recruited dynein (Fig 3K), and that these pulling forces are the primary determinant of spindle rotational mechanics.

### Cortical proteins tune dynein patterning, astral microtubule forces, and the mechanics of spindle rotation

We next investigated how other proteins influence the mechanics of spindle rotation. We performed RNAi on nine different proteins previously shown to have a role in spindle positioning ^13,24,26,28,30,31,55–60^ – *gα1/3*, *gα2*, *GTSE1*, *MARK2*, *DLG1*, *MCAK*, *TUBGCP4*, *DYNC1H1*, and *KIF13B* – and used magnetic tweezers to measure the spindle’s rotational drag coefficient. RNAi of *DYNC1H1* prevented the formation of a bipolar spindle, making magnetic tweezer experiments difficult to interpret. Of the rest of the RNAi conditions, the two proteins with the most significant impact were DLG1 and MARK2: RNAi of *DLG1* decreased the spindle’s rotational drag by more than a factor of three to 4.5 ± 1.5 pN min μm / degree (n=12, p=0.033) (Fig 4A), while RNAi of *MARK2* increased the spindle’s rotational drag by more than a factor of two to 38.0 ± 9.9 pN min μm / degree (n=9, p=0.007) (Fig 4A). While individual RNAis of the other tested proteins did not significantly impact the spindle’s rotational drag (Fig S3D, *Methods*), simultaneous RNAi of all three gα proteins (hereafter together referred to as gα) more than halved the spindle’s resistance to rotation (Fig 4A, 6.9 ± 1.8 pN min μm / degree, n=8, p=0.035). Thus, RNAi of *gα*, *DLG1*, and *MARK2* all significantly perturb the mechanics of spindle rotation. After ablation of astral microtubules, *gα, DLG1,* and *MARK2* siRNA-treated spindles did not significantly rotate, indicating a disruption of pulling forces from cortical force generators (Fig 4B).

**Fig 4.**
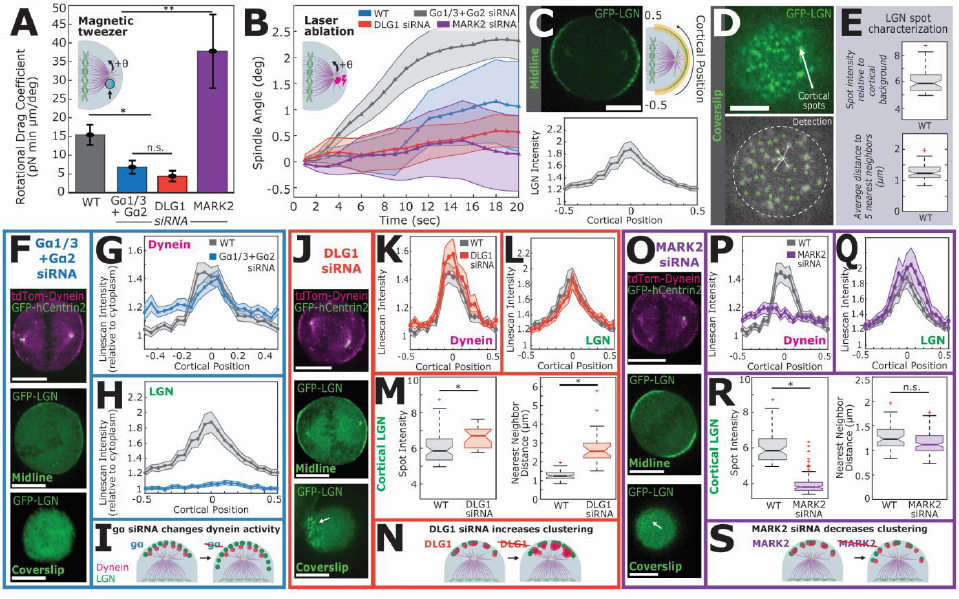
Changing the amount and extent of clustering of dynein at the cortex changes the spindle’s resistance to rotation. (A) siRNA treatments significantly changed the spindle’s resistance to rotation. RNAi of *gα* decreased the spindle’s rotational drag (n=8, p=0.035), as did RNAi of *DLG1* (n=12, p=0.033). *MARK2* siRNA treatment significantly increased the spindle’s rotational drag (n=9, p=0.007). (B) All siRNA conditions showed decreased rotation after astral microtubule ablation compared to WT (WT n = 31, gα n=10, DLG1 n=13, MARK2 n=18). (C) Cortical intensity line scans of WT U2OS cells expressing GFP-LGN show a midline crescent phenotype (n=56) with an integrated LGN intensity I_LGN_ of 16.06 ± 2.09. Scale bars in figure are 10 μm unless otherwise noted. (D) Imaging of the cortical coverslip plane of WT U2OS cells expressing GFP-LGN shows spots of LGN. Scale bar is 5 μm. (E) We quantified the intensity of LGN cortical spots relative to background, and the average distance to the five closest neighbors. (F) Cortical dynein and LGN signals in U2OS cells treated with *gα* siRNA. (G) *gα* RNAi did not affect cortical dynein localization or intensity (I_DYN_ = 7.44 ± 0.9, p=0.087 (n=26). (H) *gα* RNAi resulted in uniform, decreased cortical distribution of LGN in the midline plane with intensity I_LGN_ = 1.12 ± 0.59, p<1e-5 (n=48). (I) These results suggest that *gα* RNAi cold affect the activation state of dynein through LGN but does not change the midline crescent phenotype. (J) Cortical dynein and LGN signals in U2OS cells treated with *DLG1* siRNA. (K) *DLG1* RNAi did not affect cortical dynein localization or intensity (I_DYN_ = 7.34 ± 1.14, p=0.15) (n=32). (L) LGN midline crescent shape and intensity (I_LGN_ = 15.05 ± 1.52, p=0.71) were not changed with *DLG1* siRNA (n=46). (M) Under *DLG1* siRNA treatment, both LGN spot intensity and the distance between spots significantly increased (p=0.013 and p<0.0001). (N) These results suggest that *DLG1* siRNA increases LGN clustering. (O) Cortical dynein and LGN signals in U2OS cells treated with *MARK2* siRNA. (P) *MARK2* RNAi resulted in uniform cortical distribution of dynein (n=70), however there was no change in cortical dynein intensity (I_DYN_ = 4.75 ± 0.62, p=0.626). (Q) Midline LGN crescents were unchanged with *MARK2* siRNA treatment (I_LGN_ = 19.48 ± 3.58, p=0.38, n=38). (R) *MARK2* siRNA treatment resulted in a decreased cortical LGN spot intensity (p<0.00001), but the distance between spots stayed relatively similar (p=0.003). (S) *MARK2* siRNA delocalized cortical dynein and decreases dynein clustering within the crescents.

Next, we investigated if perturbing these proteins also impacted the patterning of LGN and dynein. In WT cells, midline cortical LGN forms crescents (Fig 4C, n=56), reminiscent of the dynein crescents (Fig 3G). Both dynein and LGN had patchy appearances in the midline crescents (Fig S4A,B), which is consistent with previous work indicating that LGN, NuMA, and dynein form clusters *in vivo* and *in vitro* ^8,19–23^. LGN also forms clearly visible spots on the cells’ cortices near the coverslip (Fig 4D), where the dynein signal was too weak to see clear patterns above background. We quantified the WT LGN spot pattern by measuring the total integrated intensity of LGN at the cortex relative to cytoplasmic intensity (WT I_LGN_=16.06 ± 2.09, n=56), the intensity of each detected spot relative to the cortical background (5.87, 95% CI: 5.62-6.11) and the average distance to the five closest neighboring spots (1.23, 95% CI: 1.16-1.30) (Fig 4E, n=28).

We compared midline dynein crescents, midline LGN crescents, and LGN spots at the coverslip across the RNAi conditions to investigate potential links between protein localization, cortical force, and spindle rotational drag. *Gα* siRNA resulted in no change in the midline dynein crescent integrated intensity (I_DYN_=7.44 ± 0.9, p=0.087) (Fig 4F,G, n=26), but produced a uniform midline distribution of LGN with significantly decreased intensity (Fig 4F,H, I_LGN_=1.12 ± 0.59, p<1e-5, n=48) and led to the disappearance of LGN cortical spots (Fig 4F). These results indicate that gα inhibition results in delocalized, uniform cortical LGN distribution, but no changes in dynein localization (Fig 4I). Interestingly, the differences between LGN and dynein signals implies that there is not a simple, linear axis between gα, LGN, and dynein, as previously suggested.

*DLG1* siRNA produced no change in the midline crescents or integrated intensity of dynein (Fig 4J,K, I_DYN_=7.34 ± 1.14, p=0.15, n=32) or LGN (Fig 4J,L, I_LGN_=15.05 ± 1.52, p=0.71, n=46), but caused the LGN spots at the cell membrane (Fig 4J) to increase in intensity to 6.69 (95% CI: 6.35-7.04, p=0.013) (Fig 4M) and become farther apart (2.57 μm, 95% CI: 2.31-2.82, p<0.0001) (Fig 4M). These results are consistent with *DLG1* siRNA causing LGN to form fewer, larger clusters without changing the total amount of cortical LGN. Further, these results point to separate regulation of clustering and midline crescent phenotypes. Given that dynein and LGN both form similar spots in crescents at the cell midline, it is reasonable to conclude that increased LGN clustering at the coverslip is indicative of increased dynein clustering (even though dynein cannot be directly quantified at the coverslip due to its low signal-to-noise there). Therefore, *DLG1* siRNA likely increases LGN and dynein clustering, while preserving their overall distribution in midline crescents (Fig 4N).

Finally, *MARK2* siRNA resulted in dynein at the midline spreading uniformly around the cortex (Fig 4O,N, n=70), while midline LGN remained in crescents (Fig 4O,Q, I_LGN_=19.48 ± 3.58, p=0.38, n=38). However, the integrated intensity of dynein at the cortex did not change from WT (I_DYN_=4.75 ± 0.62, p=0.626), indicating that a similar amount of dynein was still present at the cortex. LGN spot intensity decreased to 3.80 (95% CI: 3.73-3.87, p<0.00001) (Fig 4R), while the distance between spots stayed relatively similar (1.11 μm, 95% CI: 1.07-1.16, p=0.003) (Fig 4R). Thus, *MARK2* RNAi treatment appears to decrease LGN clustering (Fig 4S). As with the *gα* and *DLG1* siRNA phenotypes, these results seem to indicate that LGN midline crescents and LGN clustering are controlled separately.

Taken together, these results show that disruption of cortical pulling forces by knockdown of gα, DLG1, and MARK2 are accompanied by changes in the patterning of dynein and LGN.

### Mathematical modeling shows that pulling from dynein, and dynein clustering, are sufficient to explain the spindle’s rotational resistance

Our experimental results indicate that: 1) the spindle’s resistance to rotation primarily comes from pulling forces on astral microtubules that contact the cell cortex (and not cytoplasmic viscosity); 2) each of these astral microtubules is pulled on by a single molecule of dynein; and 3) the cortical force generation machinery, LGN and dynein, form “spots” on the cell cortex, and molecular perturbations that impact the spindles resistance to rotation also impact this patterning. To further investigate how cortical pulling forces could govern the resistance to rotation of the spindle, and how that could be impacted by the patterning of LGN and dynein, we next developed a three-dimensional mean field theory based on interactions between microtubules and cortically localized force generators ^45^ (*Supplementary Information*). In this model, a spindle of length 2*l*, is in a cell of radius *R*, with *M* cortical force generators distributed over the cell surface (Fig 5A). Microtubules are nucleated at the two ends of the spindle (i.e. the centrosomes) at rate *γ*, grow with speed *V_g_*, and catastrophe with rate λ (Fig 5B). A microtubule that contacts an unoccupied force generator binds to it, where upon it is subject to a pulling force of magnitude *f*_0_ directed along its length until it unbinds, which it does with rate *κ* (Fig 5C). We assume that each force generator can only bind to one microtubule at a time and, consistent with our experimental findings, that each microtubule can only bind to a single force generator at a time. Previous theoretical work has shown that such a stoichiometric interaction between microtubules and cortical force generators allows the spindle to be stably positioned with only pulling forces, which would be destabilizing if there was no limit on the number of microtubules each cortical force generators could bind ^47^. We model the observed cortical “spots” as clusters, each containing *N* individual force generators. In this model, clustering is only a statement about physical proximity of the force generators (i.e. clustering is not associated with any change in the biochemical activities of the force generators): if a microtubule contacts the cortex within the capture radius *r* of a cluster with multiple unoccupied force generator, it will bind to one of the unoccupied force generators at random.

**Fig 5.**
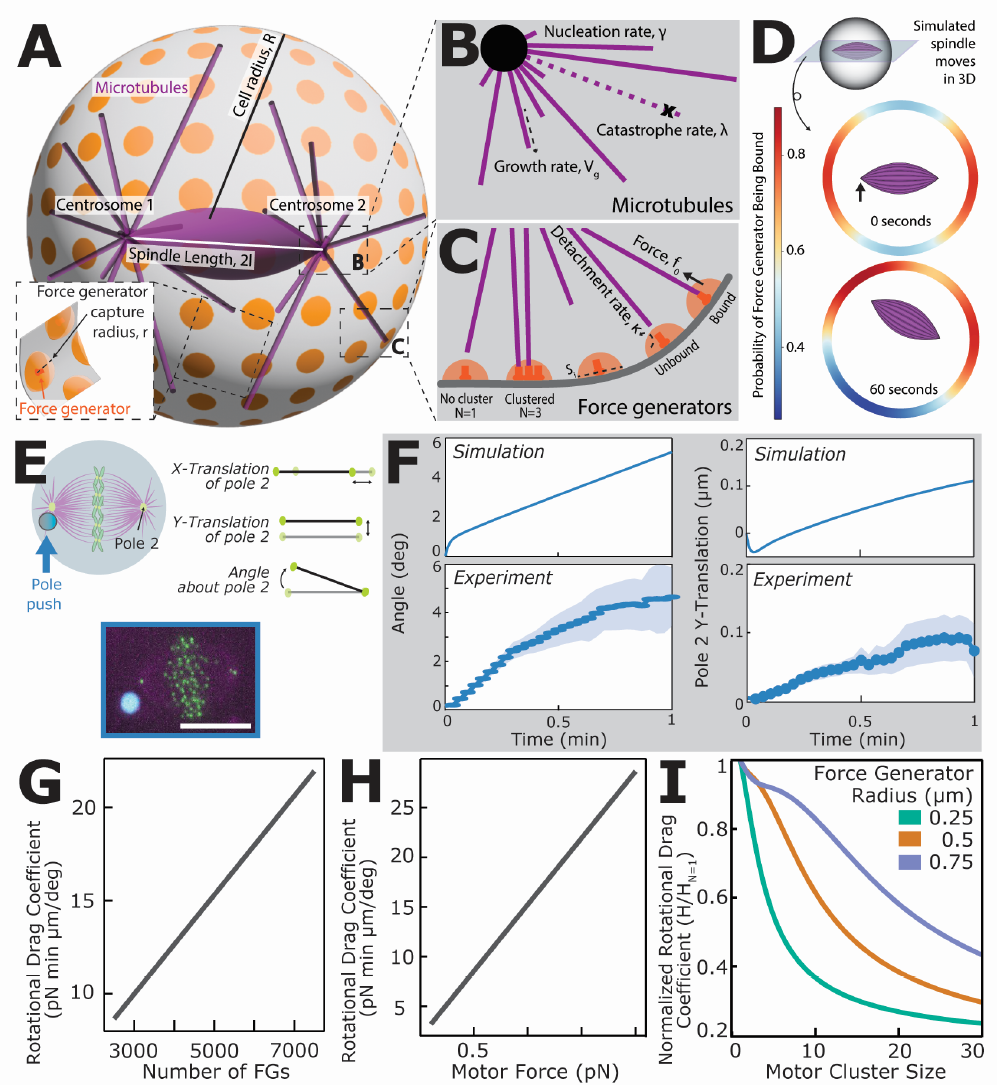
A three-dimensional mean field shows that changes in dynein clustering are sufficient to explain changes in the spindle’s resistance to rotation. (A) Overview of geometry and variables for mean field theory showing a cell with radius *R*, containing microtubules (purple), force generators (orange), and the two centrosomes. (See *Supplementary Information*) (B) Microtubules nucleate from each centrosome with rate γ, grow at rate *V_g_*, and spontaneously depolymerize with rate *λ*. (C) Force generators are located at the cell cortex, each covering an area *S_i_* equal to the intersection of the cell boundary with a sphere of radius *r*. Force generators exert force *f*_0_ on bound microtubules, and microtubules can detach with rate *κ*. (D) The probability of a force generator being bound is calculated for the entire surface of the spherical cortex. A midline slice at t = 0 seconds (top) shows a symmetrical probability distribution with motors most likely to be bound around the spindle poles. After a 60 second push (bottom, push location denoted by black arrow), the bound motor probability distribution shifts as the spindle moves closer to one part of the cortex. (E) Experimental setup to compare spindle translation and rotation. (F) The theory quantitatively matches the experimental results where pushing at the pole results in a large rotational response and very little translation (n=26). (G) Increasing the number of cortical force generators results in a linear increase of the spindle’s rotational resistance. (H) Increasing the force exerted by individual force generators also results in a linear increase of the spindle’s rotational resistance. (I) Increasing motor cluster size results in a non-linear decrease in the spindle’s rotational resistance, consistent with experimental data. Smaller individual force generators result in larger effects from clustering.

We next performed 3D simulations of this model using reasonable parameters based on our measurements or estimates (*Supplementary Information*). Simulating the application of an external force to one of the spindle poles causes the spindle to displace and rotate, which leads to variations of the attachment of cortical force generations to astral microtubules, and thus, variations in the forces acting on the spindle (Fig 5D). The resulting motion of the spindle was very similar to what we observed in the magnetic tweezer experiments (Fig 5E): In both simulations and experiments, pushing on one pole caused the spindle to rotate around the opposite spindle pole, with very little translation (Fig 5F). Thus, the model quantitatively recapitulates the rotational and translational responses of the spindle.

We next investigated how cortical force generators impact the spindle’s rotational drag coefficient (*Supplementary Information*). Increasing either the number of cortical force generators, *M*, or the magnitude of the force exerted by each force generator, *f*_0_, caused a linear increase in the simulated rotational drag coefficient (Fig 5G,H). In contrast, increasing the degree of clustering, *N*, caused a non-linear decrease in the rotational drag coefficient (Fig 5I). The extent that increasing clustering decreases the rotational drag coefficient depends on other parameters, with, for example, force generators with a smaller capture radius, *r*, exhibiting a more dramatic decrease (Fig 5I, Fig S5). While an increase in rotational drag coefficient with more force generators or increasing force per force generator is intuitive, the reason that clustering decreases the rotational drag coefficient is less obvious. To gain further insight, we analyzed the model in more detail (*Supplementary Information*). In this model, the total rotational drag of the spindle, *H_tot_*, is a sum of two terms, the rotational drag due to the cytoplasm, *H_cyto_*, and the rotational drag due to the interaction of astral microtubules with cortical force generators, *H_CFG_ : H_tot_* = *H_cyto_ + H_CFG_*. We were able to derive an analytical expression for the rotational drag coefficient due to the cortical force generators as

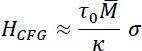

where τ_0_ = *lf*_0_ is the torque applied to the spindle by a single force generator, *M̄* = *P̄M* is the average number of cortical force generators bound to astral microtubules, *P̄* is the average probability that a cortical force generator is bound to an astral microtubule, and *σ* is a dimensionless geometric factor that depends on microtubule dynamics and cell size. Thus, in this model, the rotational drag coefficient due to cortical force generators is roughly the total torque that cortical force generators exert on the spindle, τ_0*M̄*_, multiplied by the typical time they are attached to astral microtubules, 1⁄κ. The impact of clustering enters through *P̄*, with *P̄* decreasing with increasing clustering as approximately 1⁄*N* due to competition between cortical force generators in the same cluster for astral microtubules (*Supplementary Information*). Thus, this calculation demonstrates that the patterning of force generators (i.e. whether they are arranged in clusters or spread diffusely on the cortex) is expected to have a large impact on the mechanics of spindle orientation, even if clustering does not alter the biochemical properties of individual cortical force generators.

In order to use the model to investigate the mechanics of spindle orientation in WT cells, we next sought to estimate the total number and cluster size of cortical force generators (i.e. cortically bound dynein). We estimated the number of cortical clusters using WT LGN spot density (Fig 4) multiplied by the average cell surface area, giving a range of ∼500 to 2000 clusters per cell. Previous studies indicate that LGN can form hexameric complexes ^8,19,20,23^, suggesting that dynein may cluster in groups of ∼6 (assuming 1:1 ratios of dynein and LGN). Taking a rough estimate of 1000 clusters with 5 motors per cluster (*N*=5) gives a total of 5000 motors (*M*=5000), and taking each bound motor to exert 1 pN of force (*f*_0_ = 1 pN; consistent with the measurements described above), produces a theory predicted rotational drag coefficient for the spindle of *H_tot_* = 15.3 pN min/ μm / degree, which is statistically indistinguishable from the experimentally measured value *H_WT_* =15.5 ± 2.7 pN min μm / degree (p = 0.06). Thus, pulling forces on astral microtubules by cortically bound dynein are sufficient to explain the mechanics of spindle rotation in WT cells.

Our microscopy results indicate that RNAi of *LGN* decreases the total amount of dynein at the cell cortex, RNAi of *MARK2* decreases clustering, and RNAi of *DLG1* increases clustering. We next sought to use our model to determine if these changes in localization and patterning are sufficient to account for the corresponding impact on the mechanics of spindle orientation. Assuming that *LGN* RNAi reduces dynein at the cortex by about a factor of 2.5 (*M_LGN,i_* = 2000), due to either a partial knockdown or multiple mechanisms of dynein recruitment to the cortex, then the predicted rotational drag coefficient becomes 7.3 pN min/ μm / degree, which is similar trend to experimentally observed reduction to 3.4 ± 0.6 pN min μm / degree (though the predicted and observed values are quantitatively different p =0.00009, Fig 6A,B). Assuming that RNAi of *MARK2* eliminates clustering (*N_MARK2,i_* = 1), without changing the amount of dynein at the cortex, results in a predicted rotational drag of 30.3 pN min μm / degree, which quantitatively matches the experimental measurement of 38.0 ± 9.9 pN min μm / degree (p=0.30, Fig 6A,B). Finally, postulating that RNAi of *DLG1* leads to a four-fold increase in clustering (*N_DLG1,i_* = 20), without changing the amount of dynein at the cortex, leads to a predicted rotational drag of 5.6 pN min μm / degree, which is in agreement with the experimentally observed value of 4.5 ± 1.5 pN min μm / degree (p=0.08, Fig 6A,B).

**Fig 6.**
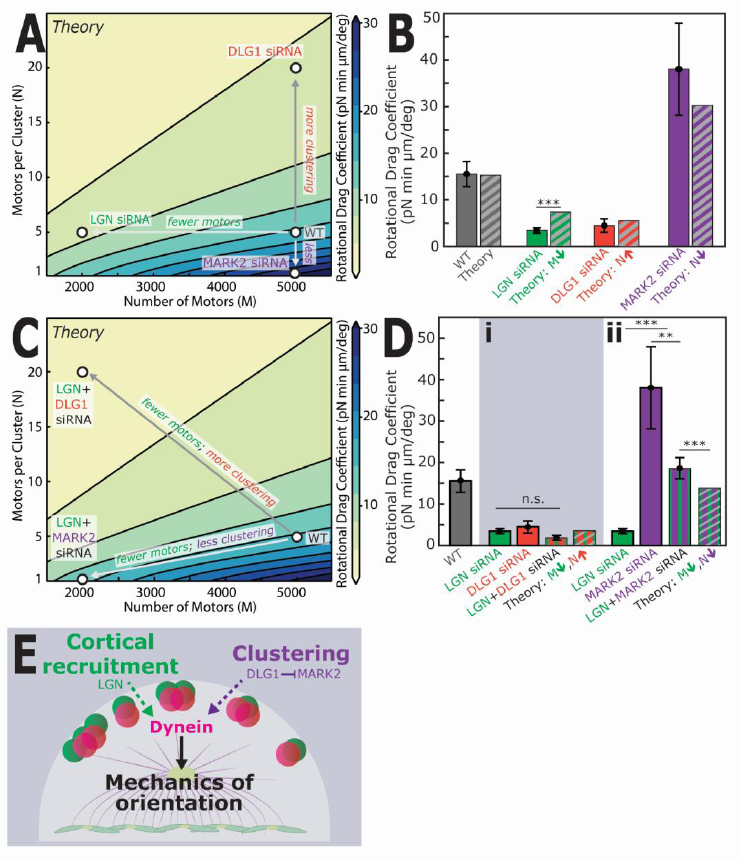
Dynein clustering and activation determine the force on the spindle. (A) Varying the number of cortical motors (M) and their clustering (N) is predicted to change the spindle’s rotational drag coefficient. By decreasing motor number from WT, the theory predicts a decrease in rotational drag (grey to green). An increase in clustering (grey to orange) results in a decreased rotational drag coefficient, while a decrease in clustering (grey to purple) results in an increased rotational drag coefficient. (B) Using M=5000 and N=5, we find quantitative agreement between the measured rotational drag and theory prediction (p=0.06). Under *LGN* siRNA treatment, we assume M drops to 2000 resulting in a predicted rotational drag coefficient of 7.3 pN min μm / degree (experiment: 3.4 ± 0.6 pN min μm / degree) (p=0.000091). Increasing motor clustering to N=20 predicts the decreased rotational drag coefficient in the *DLG1* siRNA condition (theory: 5.6 pN min μm / degree, experiment: 4.5 ± 1.5 pN min μm / degree; p=0.08). If *MARK2* siRNA results in decreased motor clustering and setting N=1, the theory predicts the resulting increased rotational drag coefficient (theory: 30.3 pN min μm / degree, experiment: 38.0 ± 9.9 pN min μm / degree; p=0.30). (C) We modeled the spindle’s response to changing M and N at the same time. (D, i) U2OS cells treated with both *LGN* and *DLG1* siRNAs showed no difference in the spindle’s rotational drag (1.8 ± 0.6 pN min μm / degree, n=17) compared with RNAi of *LGN* or *DLG1* alone (p=0.22 and p=0.11, respectively). All 3 conditions had rotational drag coefficients significantly decreased compared to WT (LGN/DLG1, p=0.01; LGN, p=0.00041; DLG1 p=0.03). This quantitatively matches the theory prediction of 3.4 pN min μm / degree, if a double siRNA treatment of *LGN* and *DLG1* would result in reduced M and increasing N (M=2000, N=20; p=0.24). (ii) Treatment with both *LGN* and *MARK2* siRNAs rescued the drag coefficient decrease caused by *LGN* RNAi (p=0.00032) and the increased caused by *MARK2* RNAi alone (p=0.011), resulting in a drag coefficient similar to WT (18.6 ± 2.6 pN min μm / degree, n=16, p=0.39). Decreasing both M and N (M=2000, N=1) results in a theoretical prediction of 13.7 pN min μm / degree, quantitatively matching our experimental results and showing the return to a WT phenotype (p=0.009). (E) Cortical force generation is regulated in multiple ways, including the cortical recruitment and clustering of dynein.

To further investigate these proposed explanations of the influence of LGN, DLG1, and MARK2 on the mechanics of spindle orientation, we next explored the impact of double knockdowns. If LGN and DLG1 independently regulate dynein localization and clustering, then the predicted impact of the double knockdown would be the combined effect of the single knockdowns described above (i.e. with *M_LGN/DLG1,i_* = *M_LGN,i_* = 2000 and *N_LGN/DLG1,i_* = *N_DLG1,i_* = 20), giving a predicted rotation drag coefficient of 3.4 pN min μm / degree (Fig 6C). We performed simultaneous RNAi of both *LGN* and *DLG1* and used magnetic tweezers to measure the spindle’s rotational drag, giving a value of 1.8 ± 0.6 pN min μm / degree, (n=17), which, within error, is consistent with the theoretical prediction (p=0.24). If LGN and MARK2 independently regulate dynein localization and clustering, then the predicted impact of the double knockdown would be the combined effect of the single knockdowns described above (i.e. with *M_LGN/MARK2,i_* = *M_LGN,i_* = 2000 and *N_LGN/MARK2,i_* = *N_MARK2,i_* = 1), giving a predicted rotation drag coefficient of 13.7 pN min μm / degree (Fig 6C), which, counterintuitively is similar to WT spindles even though both single knockdowns are different. We performed simultaneous RNAi of both LGN and MARK2 and measured the spindle’s resistance to rotation to be 18.6 ± 2.6 pN min μm / degree (n=16), which is quite similar to (but quantitatively different from, p=0.009) the predicted value. Thus, the model successfully predicts the results of both *LGN/DLG1* double knockdown and *LGN/MARK2* double knockdown, which argues both for the validity of the model and for the contention that LGN impacts the mechanics of spindle rotation by regulating the amount of dynein at the cortex, while the impact of DLG1 and MARK2 is based on their regulation of dynein clustering.

Next, we measured the spindle’s rotational drag coefficient in a *DLG1/MARK2* double siRNA condition and found that the rotational drag coefficient in the double siRNA condition (1.7 ± 0.5 pN min μm / degree, n=25) decreased relative to both WT (p=0.006) and *MARK2* siRNA (p=0.004), but was not significantly different from the *DLG1* RNAi condition alone (p=0.06) (Fig S3H). These results suggest that the MARK2 phenotype requires DLG1 expression. Given our previous results that MARK2 and DLG1 have opposite effects on clustering, it could be that MARK2 plays a regulatory role for DLG1, which is then a downstream regulator of clustering. This relationship is consistent with previous reports that MARK2 phosphorylates and inactivates DLG1 ^61^.

## Discussion

Here, we used direct force measurements and targeted molecular perturbations to investigate the mechanics of spindle orientation in human cells. We showed that the mechanics of spindle orientation are primarily determined by cortical dynein exerting pulling forces on astral microtubules and are highly sensitive to changes in the clustering of cortical dynein.

Since subcellular structures such as the spindle are small, move at slow speeds, and are surrounded by the viscous cytoplasm, they exist in the low Reynolds number regime in which inertial forces are negligible relative to viscous forces ^40^. Thus, viscous forces from the cytoplasm are a key component of most biophysical models of subcellular mechanics and dynamics. Consistent with this expectation of the importance of cytoplasm rheology, recent experiments in sea urchin oocytes indicate that the viscoelastic of the cytoplasm is a major contributor to mitotic spindle positioning ^38,62^. In contrast, we found that in human tissue culture cells, the spindle’s rotational drag coefficient is much higher than expected solely from cytoplasmic viscosity and is not impacted when cytoplasmic viscosity is changed, arguing that cytoplasmic viscosity does not significantly contribute to the mechanics of spindle orientation in human mitotic cells. One possible explanation for this difference is that, in larger cells such as in sea urchin, *Xenopus*, and zebrafish embryos ^38,62–64^, astral microtubules are not long enough to contact the cortex so the mechanics of the cytoplasm significantly impact spindle positioning and orientation, while in smaller cells, such as human tissue culture cells and in *C. elegans* embryos ^36,45^, viscous forces are negligible because the interactions between astral microtubules and the cell cortex dominate over all other forces.

The nature of the interactions between astral microtubules and the cortex have been controversial ^8,9,65^. Our observation that cutting astral microtubules with a laser causes the spindle to rotate away from the cut demonstrates that astral microtubules in human tissue culture cells are subject to net pulling forces. Removing dynein from the cortex using *LGN* siRNA causes a dramatic decrease in both pulling forces (revealed via laser ablation) and the spindles’ resistance to rotation (revealed via magnetic tweezers), indicating that the pulling forces on astral microtubules are primarily generated by cortical dynein and that these pulling forces are a major determinant of the mechanics of spindle orientation. Using the speed that the spindle rotates when astral microtubules are cut (via laser ablation), the force required to cause this motion (measured via magnetic tweezers), and measurements of the number of cut astral microtubules (via EB1 imaging), we calculate that each astral microtubule that contacts the cortex is subject to ∼1 pN of pulling force, consistent with each astral microtubule being pulled on by a single molecule of dynein ^49–54^. We construct a biophysical model of the consequences of pulling forces from cortical dynein and find that they are sufficient to explain the measured resistance to rotation of the spindle. Thus, while other forces – such as viscous forces, microtubules pushing and bending forces, and pulling forces from microtubule depolymerization – may be present, our results argue that they do not significantly contribute to the mechanics of spindle orientation in human mitotic cells, which primarily results from pulling forces from cortically anchored dynein.

Previous work has shown that many molecules regulate the behavior of dynein on the cortex, including determining how dynein clusters, and that this behavior is important for spindle positioning. Light microscopy indicates that dynein and associated proteins form clusters on the cell cortex in human cells ^20^ and yeast ^22,66–68^. *In vitro* studies show that LGN and NuMA co-oligomerize, which has been proposed to be the basis of dynein clustering *in vivo* ^19,23^. Even dynactin, which is thought to activate dynein, may play a role in clustering by cross-linking two dynein molecules together ^69^. Preventing the oligomerization of LGN or NuMA (and thus presumably preventing dynein clustering) results in misoriented spindles in both human cells ^20^ and yeast ^68^. Over-clustering dynein artificially in human cells also causes spindle orientation defects, including increased spindle rocking ^8,20^. Thus, dynein clustering seems to be important for spindle positioning and orientation, but the mechanism remains unclear.

Our biophysical modeling indicates that clustering has a major impact on spindle positioning and orientation solely by virtue of co-localizing dyneins, without needing to invoke any changes to the biochemical or biophysical properties of dynein. In our model, motor clustering leads to competition for free astral microtubule ends that contact the cortex in a given location, which means that for a fixed number of force generators, as clustering increases, the probability of a given force generator being bound to a microtubule decreases. A cluster can be seen as an intermediate between stoichiometric binding (i.e. one force generator per astral microtubule, in which pulling forces can be stabilizing) and non-stoichiometric binding (i.e. all astral microtubules that contact a force generator can bind, in which pulling forces are always destabilizing) ^47^, meaning that as the number of motors in a cluster increases, the spindle’s orientation and position become more unstable.

We showed that DLG1 and MARK2 both impact clustering (via light microscopy of siRNA-treated cells) and the mechanics of spindle orientation (via magnetic tweezers). Our modeling indicates that the impact of DLG1 and MARK2 on clustering is sufficient to explain their impact on the mechanics of spindle orientation. Removing *DLG1* by siRNA treatment resulted in a decreased rotational drag coefficient, increased LGN spot intensity without changing the total amount of cortical dynein, and increased distance between LGN spots, consistent with DLG1 decreasing LGN and dynein clustering. Removing *MARK2* by siRNA treatment increased the spindle’s rotational drag and decreased LGN spot intensity without changing the total amount of cortical dynein, suggesting that MARK2 increases LGN and dynein clustering. Further, the simultaneous RNAi of *MARK2* and *DLG1* is similar to wildtype, indicating that MARK2 requires DLG1 in order to impact clustering. MARK2 has been previously studied in neurons, where it has been shown to phosphorylate DLG1, causing DLG1 to lose its ability to properly localize to the synaptic membrane, where it normally acts as a scaffold protein^61^. We hypothesize that in mitotic human cells, MARK2 also acts by phosphorylating DLG1, thereby inhibiting DLG1’s ability to suppress LGN and dynein clustering. Neuronal DLG1 is also regulated by cdk5-dependent phosphorylation, which inhibits DLG1’s oligomerization at the synapse ^70^. The role of DLG1 in neurons as an oligomerizing scaffold protein negatively regulated by phosphorylation is similar to its proposed behavior involving dynein at the cell cortex in our model.

Further, studies have shown that the presence of DLG1 and its direct interactions with LGN are required for proper spindle orientation and that LGN signal becomes patchier on both the cortex and in the cytoplasm with *DLG1* siRNA ^24,26^, suggesting that DLG1 may have a role in recruiting and organizing LGN at the cortex. Some studies have suggested that DLG1 anchors a separate, parallel force generation pathway involving KIF13B ^57,71,72^ that may also help to load astral microtubules onto cortical dynein, however we saw no difference in the spindle’s rotational drag after *KIF13B* RNAi, indicating that this pathway does not significantly contribute to cortical force generation. Therefore, we propose that DLG1 binds to LGN, which stabilizes LGN at the cortex and prevents it from oligomerizing (or potentially co-oligomerizing with NuMA) and thus decreases dynein clustering. MARK2 prevents this inhibition by phosphorylating DLG1.

Our experimental results and modeling show that both dynein recruitment to the cortex and dynein’s cluster state once at the cortex are important for the mechanics of spindle orientation. We propose that different molecular pathways regulate these different processes: i.e. that LGN (and associated factors) recruit dynein to the cell cortex, while DLG1 (and associated factors) determines dynein’s cluster state once it has reached the cortex. Previous studies have reported varying results when recruiting dynein to the cortex artificially. In some cases, authors argue that simply recruiting dynein is sufficient for successful spindle orientation ^16^, while in other studies, targeting dynein to the cortex resulted in misoriented spindles with increased spindle rocking ^20^. One potential explanation for these different results is that the impact of recruiting dynein to the cortex depends on the cluster state of dynein once it reaches the cortex. Okumura *et al*. ^20^ reported that using their optogenetic system to recruit dynein to the cortex resulted in a much larger amount of cortical dynein than would be present in wildtype cells, potentially mirroring a state of increased dynein clustering, which could explain why their method of dynein recruitment resulted in misoriented, unstable spindles.

Cluster size is a relatively easy molecular property to quickly change. In the context of mitosis where dynamic regulation of forces is important for successful division, our proposed mechanism suggests that increasing or decreasing stability of the spindle would not require any protein breakdown nor would it require extensive remodeling of the astral microtubule array. Instead, changing the cluster state of dynein at the cortex could quickly result in a stiffer or more mobile spindle orientation. This control mechanism of using cluster state to influence force generation could be more broadly relevant in the cell. For example, dyneins cluster *in vivo* to collectively move phagosomes of varying sizes ^73^. Evidence of clustering has also been seen in other motors, such as when kinesin clustering enables continuous vesicle towing along microtubules, which could not be achieved by a single kinesin ^74^. Although the mechanism by which clustering may impact force generation in these other systems has not been determined, the functional importance of clustering points towards the idea that clustering could be a broad control mechanism used in the cell.

## Acknowledgements

The authors would like to thank Xingbo Yang, Kuan-Chung Su, and Iain Cheeseman for valuable discussions. This work was funded by NSF award DBI-1919834. M.A.-D. was supported by the QuantBio Student Fellowship through the NSF-Simons Center for Mathematical and Statistical Analysis of Biology at Harvard (#1764269). S.B. was supported by NIH R01 (GM131004) and NSF CAREER 1846010. H.M was supported by NIH R01 (GM131004) to S.B., the Gruber Foundation, and NIH T32s (GM100884 and GM007499). We also acknowledge support from the CCBX program of the Center for Computational Biology of the Flatiron Institute.

## Author Contributions

M.A.-D. and D.J.N. designed the study. M.A.-D. performed magnetic tweezer experiments, siRNA experiments and imaging, and EB1 imaging. M.A.-D. and H.W. performed laser ablation experiments. V.G.H., R.F., and M.J.S. derived the mean field model and wrote the theory supplement with input from M.A.-D. and D.J.N. M.A.-D. performed data processing and statistical analyses. H.M. created the CTDNEP1-KO cell line and was supervised by S.B. V.G.H. was supervised by R.F. and M.J.S. M.A.-D. and H.W. were supervised by D.J.N. M.A.-D. and D.J.N. wrote the paper with input from all authors.

## Methods

### Cell culture

U2OS cell lines were maintained in Dulbecco’s Modified Eagle Medium (DMEM, Gibco) supplemented with 10% Fetal Bovine Serum (FBS, Gibco), 50 IU/mL penicillin, and 50 μg/mL streptomycin (both Gibco) at 37 °C in a humidified atmosphere with 5% CO_2_. Cells were passaged every 2-3 days. Several U2OS lines stably expressing fluorescent fusion proteins were used: U2OS with GFP-hCentrin2, mCherry-alpha tubulin, and GFP-CENPA; U2OS with GFP-hCentrin2 and tdTomato-DYNHC2; U2OS ΔCTDNEP1 as described in ^44^ with GFP-hCentrin2; U2OS with GFP-LGN. Cells were validated as mycoplasma free by a PCR-based mycoplasma detection kit (Sigma Aldrich).

### Magnetic tweezer system

The cylindrical magnetic tweezer core was made of ¼” wide, 6” long HyMu80 alloy (EFI Alloy 79, Ed Fagan Inc.) and sharpened at one end to a cone with a tip width of 5 μm. The solenoid frame was a steel cylinder 1” wide, 3” long, and had a hole for the core to fit through. The frame was held onto the core using 2 set screws. The solenoid was made using sheathed, 24-gauge copper wire (7588K77, McMaster-Carr) wound 400 times. The solenoid and core were mounted on a micromanipulator (NMN-21, Narishige), which was mounted on a custom-built base using ThorLabs components. The solenoid was connected to a programmable power supply (PSP-603, GW Instek). (See system diagram in Fig S1A.) To calibrate the system, we tracked superparamagnetic beads moving through pure glycerol due to force produced at the tweezer tip by a set current running through the solenoid. We used these trajectories to interpolate a 2D force map around the tweezer tip including 95% confidence interval surfaces (Fig S1C,D). After interpolation, we compared the calibration trajectories (with known velocity and force) to the values that would be extracted from the calibration map to check the calibration map (Fig S1E). During bead push experiments, we tracked the positions of the tweezer tip and bead and then used that relative trajectory information to determine instantaneous force on the bead throughout the push (see “Quantitative Image Analysis” below).

### Fluorescence imaging

Bead push experiments, laser ablation experiments, and all fluorescence imaging were performed on a Nikon Eclipse Ti microscope equipped with either a manual rotation stage (bead push experiments) or a motorized XY stage (laser ablation experiments, fluorescence imaging). Multi-dimensional time-series images were acquired with MicroManager controlling a Yokogawa CSU-X1 spinning disk unit with a 1.2X camera mount magnifier, Coherent Obis lasers (488, 560, 640 nm), a motor-driven filter wheel (96A353, Ludl Electronics Products with filters 514/30 BP, 593/40 BP, 647 LP), an sCMOS camera (Flash LT+, Hamamatsu), an objective z-piezo stage (P-721, Physik Instrumente), and an oil-immersion objective (CFI Plan Apo Lambda 60X Oil, NA 1.4, Nikon). Only metaphase cells displaying proper chromosome alignment, no significant blebbing or morphological issues, and a high expression level of the expected tags were selected for experiments.

### Magnetic tweezer experiments

Cells were grown in a 35 mm diameter, Desag 263 glass coverslip-bottomed culture dish (0.17 mm thick glass black DeltaT, Bioptechs) to 75-80% confluency. Prior to experiments, cell medium was switched to 500 μL imaging medium (FluoroBrite DMEM, Gibco) supplemented with 10 mM HEPES and left to equilibrate for 10-20 minutes. Then a layer of 1 mL white mineral oil (VWR) was added on top of the imaging medium, and the dishes were mounted in a temperature control system and kept at 37 °C (DeltaT Culture Dish System, Bioptechs). Only cells containing exactly one bead were used for all measurements. Cells were positioned relative to the fixed tweezer tip using the rotation stage. In some cases, beads were moved into position using the tweezer before pushing on the spindle. During a force experiment, the power supply was set to a stable voltage based on prior calibration measurements and turned on and off during imaging. For these experiments, we imaged 3 z-planes spaced 2 μm apart every 2 s. In some cells, there was a short delay of up to 10 s before the spindle began to rotate; this may be due to the bead moving a short distance before contacting the spindle. When averaging traces, we set t = 0 s as the beginning of rotation.

### siRNA

Cells were grown to ∼70% confluency in DMEM/10% FBS/penstrep as described above. For each dish, an siRNA transfection cocktail was prepared. First, 300 μL OptiMEM I Reduced-Serum Medium, GlutaMAX Supplement (ThermoFischer) and 3.2 μL of 100 μM siRNA were mixed and left to incubate for 5 minutes, flicking the tube occasionally (see below for all siRNAs used). Concurrently, 300 μL OptiMEM and 2 μL Lipofectamine RNAiMAX (ThermoFischer) were mixed and left to incubate for 5 minutes, flicking the tube occasionally. Next, the siRNA/OptiMEM mix was added to the Lipofectamine/OptiMEM mixture and left to sit for 30 minutes, flicking the tube every 10 minutes. Cells were washed twice with warm PBS and then 2 mL DMEM/10% FBS (no penstrep) was added to each dish. The siRNA cocktail (∼600 μL total) was added to each dish, and cells were incubated for 24 hours. Then the cells were washed twice with PBS and split onto Bioptechs DeltaT culture dishes with DMEM/10% FBS/penstrep. Then, cells were incubated for 24 hours before use in experiments.

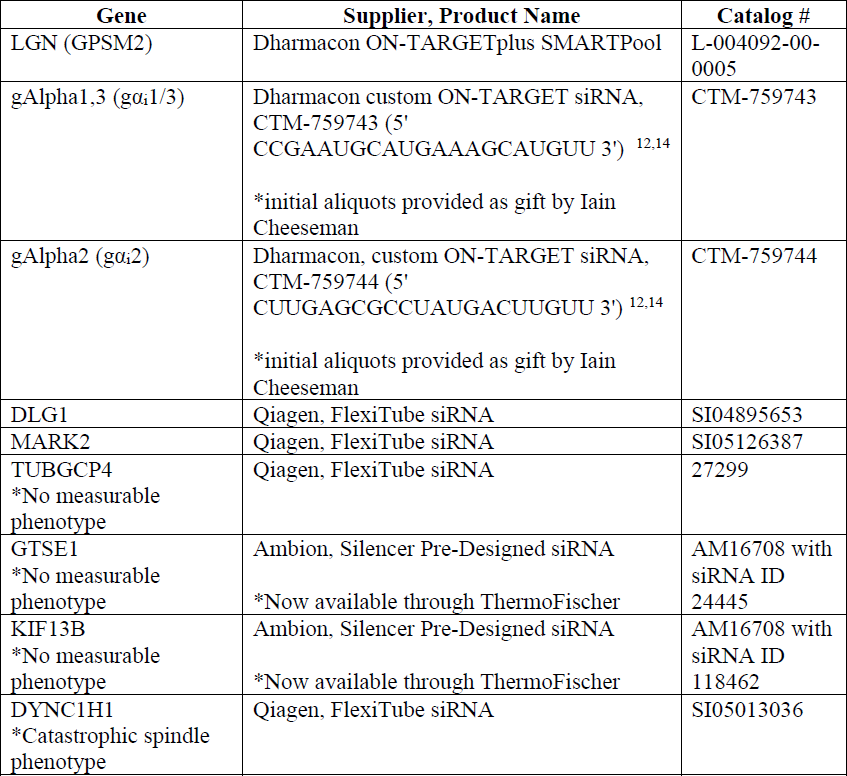

### Laser ablation

Cells were prepared as for magnetic tweezer experiments. The laser ablation system was constructed on the same spinning disc confocal microscope used for magnetic tweezer experiments and fluorescence imaging, as described previously in ^45,46^. Briefly, femtosecond pulses (80 MHz) with 0.3-nJ pulse energy and 800-nm center wavelength came directly from an Insight pulsed laser (InsightX; Spectra-Physics, Mountain View, CA). A 16-kHz femtosecond pulse train with ∼6-nJ pulse energy was produced by selecting pulses using a pulse picker (Eclipse Pulse Picker; KMLabs). The ablation laser was focused through the same objective used for imaging, and laser ablation was performed by moving the sample on a piezo-stage (P-545 PInano XYZ; Physik Instrumente) in three dimensions controlled by a home-developed LabVIEW program (National Instruments). To ablate astral microtubules, we generated a curved plane centered on the target centrosome with a radius of 2 μm, height of 5 μm, and subtending 0-60° (Fig 3A). The moving speed of the stage was 100 µm/s. After ablation, we imaged 3 z-planes approximately every second for about 60 seconds (exact frame rate varied slightly day to day based on z-piezo temperature and camera read out speeds but was recorded for each stack individually).

### EB1 measurements

We transfected WT U2OS cells with EB1-GFP (1 µL Lipofectamine 2000, ThermoFischer, with 100 ng plasmid) and incubated cells for 24 hours. The next day, cells were split onto Bioptechs DeltaT culture dishes with DMEM/10% FBS/penstrep and incubated for 24 hours before experiments. During the experiments, we imaged cells both at the midline (205 ms frame rate, 200 ms 488 laser exposure per frame) and coverslip (305 ms frame rate, 300 ms 488 laser exposure).

### Quantitative image analysis

*(Magnetic tweezer experiments)* A MATLAB program, based on a particle tracking algorithm ^75^, was developed to track the trajectories of kinetochores, poles, and beads within cells in 3D. Next, relative distances and angles were calculated between all points of interest. For each cell, the pole movement was divided by the tweezer force for normalization. Each movement trace in a given condition was then multiplied by the average force across all cells. We used several open-source MATLAB packages: featuretrack by Maria Kilfoil, John Crocker, and Eric Weeks; quiverc by Bertrand Dano; and shadedErrorBar by Rob Campbell. *(Laser ablation)* A modified version of the pipeline described for bead push experiments was used to track poles during laser ablation experiments and combine data between magnetic tweezer experiments (spindle lengths, rotational resistance) and laser ablation experiments (spindle rotational velocity). *(EB1)* A MATLAB program was developed to identify EB1 trajectories in both midline and coverslip images. For the midline analysis, only trajectories that were within 0.5 μm of the cortex at any point were considered “in contact”. For the cortex analysis, any comet visible at the coverslip was considered in contact with the cortex. From the midline data, we extracted the number of microtubules per cortical area (MTs/µm^2^), average cortical dwell time (s), and growth velocity (µm/s). From the coverslip data, we extracted the number of microtubules per cortical area (MTs/µm^2^) and the average cortical dwell time (s). To estimate the total number of microtubules in contact with the cortex at any given time, we multiplied the density of cortical microtubules with the total surface area of the cell. There was good agreement between midline and coverslip data, and we report an estimated range based on both methods. *(Dynein and LGN midline patterning)* A MATLAB program was developed to identify the cell boundary and calculate a line scan of cortical fluorescence intensity for either GFP-LGN or tdTomato-dynein signal. For each cell, all intensities were first normalized to background intensity outside the cell far from the cortex. Then the cortical intensity was divided by the average cytoplasmic fluorescence signal, and the perimeter was split into 2 half-cell traces, using the cortical coordinates intersecting with the metaphase plate. Then, the half-cell perimeter was scaled to a dimensionless cortical coordinate (-0.5 to 0.5, with 0 being at the pole). After normalizing all intensity scans, each line scan was multiplied by the average cytoplasmic fluorescence intensity across all cells. *(LGN coverslip patterning)* A MATLAB program, based on featuretrack by Maria Kilfoil, was developed to identify LGN spots. We extracted spot intensity normalized to cortical background and calculated the distances between each spot’s 5 nearest neighbors. Due to high variability in cortical patterning, we report the median, lower quartile, and upper quartile values of the data.

### Statistical analysis

Statistics are presented as mean ± SEM (unless otherwise noted), and *p* values were calculated using either one-sample (comparison of experiment to theory) or paired (comparison between experimental conditions) t-tests (ttest, ttest2 in MATLAB, respectively).

