## Supplementary Information for "Clustering of cortical dynein regulates the mechanics of spindle orientation in human mitotic cells"

**A theoretical estimate of the rotational drag coefficient of the spindle due to interactions with the cytoplasm.**

We approximate the spindle as a prolate ellipsoid, having a major axis length $2l$ and minor axes length $2l_s$, rotating in a viscous fluid [1,2]. We set $l = 5$ µm and $l_s = 1.5$ µm, and define the effective spindle radius as $\rho = \frac{l}{l_s} = 3.33$. The rotational drag coefficient for the ellipsoid about a short axis is given by

$$H_{cyto} = 8\pi\eta\bar{l}^3\alpha \qquad (1)$$

where $\bar{l} = (l l_s^2)^{\frac{1}{3}}$ , $\eta$ is the cytoplasmic viscosity, and the shape parameter $\alpha$ is given by

$$\alpha = \frac{2}{3\rho}\frac{(\rho^2+1)(\rho^2-1)^{\frac{3}{2}}}{(2\rho^2-1)\ln\left(\rho+\sqrt{\rho^2-1}\right)-\rho\sqrt{\rho^2-1}} \qquad (2)$$

This estimation assumes that the spindle rotates about its center. We multiply $\alpha$ by a factor of two to account for the experimental observation that the spindle rotates around a pole.

**A 3D mean field cortical force model.**

***The set up.***

We present a generalization of the independent force generator model developed in [3]. Consider a spindle of fixed length $2l$, with its centrosomes at positions $\boldsymbol{x}_k(t)$ ($k = 1, 2$; $|\boldsymbol{x}_1 - \boldsymbol{x}_2| = 2l$) at time $t$, moving inside a spherical cell of radius $R$ whose interior surface holds an assembly of force generators (Fig 5A, Table S1). We assume that microtubules nucleate uniformly in all directions from each centrosome at rate $\gamma$, with each growing at speed $V_g$ and randomly going through catastrophe at rate $\lambda$ and immediately shrinking (Fig 5B, Table S1). In the absence of a cell boundary, the average length of microtubules emanating from each centrosome is $L = V_g/\lambda$. Microtubules that reach the cell boundary either bind to an unbound force generator or undergo catastrophe and shrink. We considered $M$ force generators divided into clusters, each containing

$N$ motors (and hence *M/N* total clusters). Each motor in a cluster shares the same capture domain of radius *r*, with their centers are at the same point. In our model, the capture domains of clusters do not overlap. We assume the interaction of microtubules and force generators is stoichiometric—each force generator can only interact with one microtubule at a time. If a microtubule binds a force generator, this interaction generates a pulling force of magnitude $f_0$ along the microtubule direction toward the motor. Bound force generators detach from microtubules with rate $\kappa$ and can then bind another microtubule. At any time, the spindle is in force balance with total pulling forces from force generators equal and opposite to drag forces from the cytoplasm. Drag forces are represented by the translational drag coefficient $\mu_{cyto}$ and the rotational drag coefficient $H_{cyto}$.

***Model description.***

Ultimately, we will derive an equation of motion for the spindle based on forces exerted by force generators bound to microtubules. First, we model the spatiotemporal density distribution of microtubule ends emanating from each centrosome. We then calculate the interaction between microtubules and force generators and use this to determine the probability that a microtubule binds to an unbound force generator. Finally, we calculate the force from force generators on centrosomes and derive the spindle's equation of motion by assuming a simple drag law.

The density distribution of microtubule endpoints nucleating from centrosome ($k = 1,2$) at time $t$, $\psi_k(\boldsymbol{x}, t)$, satisfies the Fokker-Planck equation

$$\partial_t \psi_k + \nabla \cdot \left( V_g \frac{\boldsymbol{x} - \boldsymbol{x}_k}{|\boldsymbol{x} - \boldsymbol{x}_k|} \psi_k \right) = -\lambda \psi_k + \gamma \delta(\boldsymbol{x} - \boldsymbol{x}_k) \qquad (3)$$

where $\boldsymbol{x}_k$ is the position of centrosome $k$. As long as $\mathrm{t} \gg \frac{\mathrm{L}}{\mathrm{V_g}}$, we can use the steady-state, where $\partial_t \psi_k = 0$ and the density distribution of microtubule endpoints is given by

$$\psi_k(\boldsymbol{x}) = \frac{\gamma}{4\pi V_g} \frac{1}{|\boldsymbol{x} - \boldsymbol{x}_k|^2} e^{-\frac{|x - x_k|}{L}} \qquad (4)$$

To calculate the pulling force generated by a force generator in the $i$-th cluster on the $k$-th centrosome, we first compute the rate, $\Omega_i^k(\boldsymbol{x}_k, \dot{\boldsymbol{x}}_k)$, at which microtubules from the centrosome impinge upon that cluster

$$\Omega_i^k(\boldsymbol{x}_k, \dot{\boldsymbol{x}}_k) = \int_{S_i} \psi_k(\boldsymbol{x}') \left[ \left( V_g \frac{\boldsymbol{x}' - \boldsymbol{x}_k}{|\boldsymbol{x}' - \boldsymbol{x}_k|} + \dot{\boldsymbol{x}}_k \right) \cdot \hat{\mathbf{n}}_{x'} \right]_+ dA(\boldsymbol{x}') \tag{5}$$

where $S_i$ is the surface area covered by the cluster and $[\bullet]_+ = max\{0,\bullet\}$ ensures that the impingement rate is always non-negative. If the size of the cluster is much smaller than the size of the cell, we can approximate the impingement rate by a midpoint approximation, which is second order in $\frac{r}{R}$, that gives

$$\Omega_i^k(\boldsymbol{x}_k, \dot{\boldsymbol{x}}_k) \approx 2\pi\chi R^2 \psi_k(\boldsymbol{y}_i) \left[ \left( V_g \frac{\boldsymbol{y}_i - \boldsymbol{x}_k}{|\boldsymbol{y}_i - \boldsymbol{x}_k|} + \dot{\boldsymbol{x}}_k \right) \cdot \hat{\boldsymbol{n}}_i \right]_+ \tag{6}$$

where $\boldsymbol{y}_i$ is the position of the $i$-th cluster, $\hat{\boldsymbol{n}}_i$ is the outward normal to the cell surface at the cluster center, and $\chi = 1 - \frac{1}{\sqrt{1+\left(\frac{r}{R}\right)^2}}$ is a geometric factor that comes from integrating $\int_{S_i} dA$. (Note that this approximation differs from the one used in previous presentations of this model [3,4].)

The probability of the $l$-th force generator in the $i$-th cluster being bound to a microtubule from the $k$-th centrosome, $P_{i,l}^k$, evolves as

$$\frac{dP_{i,l}^k}{dt} = -\kappa P_{i,l}^k + \Omega_i^k(\boldsymbol{x}_k, \dot{\boldsymbol{x}}_k) \mathbb{P}\left(\text{new MT binds to FG}_{i,l}\right) \tag{7}$$

where we used the notation FG for force generator.

To determine the probability $\mathbb{P}\left(\text{new MT binds to FG}_{i,l}\right)$, we make two assumptions of the interaction between force generators and microtubules. First, while a force generator is bound to a microtubule, it will not interact with any other microtubules. This is the stoichiometric assumption. Second, when a microtubule reaches a cluster, we assume that it will randomly attach to any unbound force generator in that cluster, meaning that there is no steric hindrance or other effect coming from overlapping force generators. Using the law of total probabilities, we write

$$\mathbb{P}\big(\text{new MT binds to FG}_{i,l}\big) \\ = \mathbb{P}\big(\text{FG}_{i,l} \text{ is unbound}\big)\mathbb{P}\big(\text{MT binds to FG}_{i,l} \mid \text{FG}_{i,l} \text{ is unbound}\big) \tag{8}$$

where we set that $\mathbb{P}\big(\text{MT binds to FG}_{i,l} \mid \text{FG}_{i,l} \text{ is bound}\big) = 0$.

From the stoichiometric model, $\mathbb{P}\big(\text{FG}_{i,l} \text{ is unbound}\big) = 1 - P_{i,l}^1 - P_{i,l}^2$, and we approximate $\mathbb{P}\big(\text{MT binds to FG}_{i,l} \mid \text{FG}_{i,l} \text{ is unbound}\big) \approx \frac{1}{1+\sum_{l\neq k}\mathbb{P}(\text{FG}_{i,k} \text{ is unbound})}$.

If there is only one force generator per cluster ($N = 1$), then $\mathbb{P}\big(\text{new MT binds to FG}_{i,l}\big) = \mathbb{P}\big(\text{FG}_{i,l} \text{ is unbound}\big)$, since $\frac{1}{1+\sum_{l\neq k}\mathbb{P}(\text{FG}_{i,k} \text{ is unbound})} = 1$. The probability of attachment takes the form of previous model [4]:

$$\frac{dP_i^k}{dt} = \Omega_i^k(\boldsymbol{x}_k, \dot{\boldsymbol{x}}_k)(1 - P_i^1 - P_i^2) - \kappa P_i^k \tag{9}$$

If $N > 1$, we assume that all force generators in each cluster have the same binding probabilities. Then we can solve for the second term (dropping the *l* index), and we find that the probability of attachment is given by

$$\frac{dP_i^k}{dt} = \Omega_i^k(\boldsymbol{x}_k, \dot{\boldsymbol{x}}_k)\frac{1 - P_i^1 - P_i^2}{1 + (N-1)(1 - P_i^1 - P_i^2)} - \kappa P_i^k \tag{10}$$

The total force from force-generators on each centrosome is given by

$$\boldsymbol{F}_k = \sum_{i=1}^{M/N} N f_0 \frac{\boldsymbol{y}_i - \boldsymbol{x}_k}{|\boldsymbol{y}_i - \boldsymbol{x}_k|} P_i^k \tag{11}$$

The mean field step is to approximate this sum by an integral, and replacing the discrete quantities $P_i^k(t)$ and $\Omega_i^k(\boldsymbol{x}_k, \dot{\boldsymbol{x}}_k)$ by functions over the sphere denoted by $P^k(\boldsymbol{y}, t)$ and $\Omega^k(\boldsymbol{y}, \boldsymbol{x}_k, \dot{\boldsymbol{x}}_k)$

$$\boldsymbol{F}_k \approx M f_0 \frac{1}{4\pi R^2} \int_{|y|=R} \frac{\boldsymbol{y} - \boldsymbol{x}_k}{|\boldsymbol{y} - \boldsymbol{x}_k|} P^k(\boldsymbol{y}, t) dA(\boldsymbol{y}) \,. \qquad (12)$$

Finally, we pose the equations of motion of the spindle due to pulling forces from force generators, drag forces from the cytoplasm, and an external force $\boldsymbol{F}_{ext}$ and torque $\boldsymbol{\tau}_{ext}$ on the spindle. The spindle center is at $\boldsymbol{x}_C = \frac{x_1 + x_2}{2}$, and we let $\boldsymbol{S} = \frac{x_1 - x_2}{2}$, which is the axis vector ($|\boldsymbol{S}| = \boldsymbol{2l}$). Letting ω be the spindle rotational velocity about $\boldsymbol{x}_C$, the equations of motion of spindle center and axis are given by

$$\mu_{cyto} \frac{d\boldsymbol{x}_C}{dt} = \boldsymbol{F}_1 + \boldsymbol{F}_2 + \boldsymbol{F}_{ext} \qquad (13)$$

$$\frac{d\boldsymbol{S}}{dt} = \boldsymbol{\omega} \times \boldsymbol{S} \qquad (14)$$

with

$$H_{cyto} \boldsymbol{\omega} = \boldsymbol{S} \times (\boldsymbol{F}_1 - \boldsymbol{F}_2) + \boldsymbol{\tau}_{ext}. \qquad (15)$$

Next, we analyze the behavior of the spindle under external torque in the presence of cortical force-generators.

To numerically evolve the system, we discretize the surface of the sphere using a Lebedev quadrature and use Runge Kutta 4 as the time marching scheme (see *Simulations* below).

***Determining the effective rotational drag coefficient on the spindle.***

Here, we calculate a rotational drag coefficient due to the spindle's interaction with cortical force generators ($H_{CFG}$) for slow rotations of the axis vector $\boldsymbol{S}$. That is, setting $\boldsymbol{x}_C(t) = (0,0,0)$, we assume $\boldsymbol{S}(t) = l(\cos(\omega t), \sin(\omega t), 0)$, so that $\boldsymbol{\omega} = (0,0,\omega)$.

The torque coming from force generator interactions is

$$\boldsymbol{\tau}_{CFG} = \boldsymbol{S} \times (\boldsymbol{F}_1 - \boldsymbol{F}_2) \qquad (16)$$

For small $\omega$ and over long times, we will show that there is a linear relationship between this torque and the angular velocity:

$$\boldsymbol{\tau}_{CFG} \approx -H_{CFG}\,\boldsymbol{\omega} = -H_{CFG}\,\omega\hat{\mathbf{z}} \tag{17}$$

From Eq. (17) with the expression for $\boldsymbol{\omega}$ in Eq. (15)

$$\left(H_{cyto} + H_{CFG}\right)\omega \approx \boldsymbol{\tau}_{\text{ext}} \cdot \hat{\mathbf{z}} \tag{18}$$

Therefore, we can define the effective rotational drag coefficient of the spindle as

$$H_{tot} = H_{cyto} + H_{CFG} \tag{19}$$

To compare with experiments, we multiply $H_{tot}$ by a factor of two, since in this calculation the center of rotation of the spindle is the center of the cell, while in experiments the spindle rotates around a pole.

In the following sections, we will show how to calculate $H_{CFG}$.

***Effective rotational drag: Rotating frame of reference***

From Eq. (17), the effective rotational drag is given by

$$H_{CFG} = -\frac{\mathrm{d}}{\mathrm{d}\omega} \lim_{t\to\infty} \boldsymbol{\tau}_{\mathrm{MT}} \cdot \hat{\mathbf{z}}\Big|_{\omega = 0} \tag{20}$$

The order of the operators in crucial. If we evaluate the derivative before taking the limit, we are essentially fixing the rotation to 0 and then leaving it in this configuration for all time.

For convenience, we change our reference frame to one that rotates with the spindle. We introduce the vectors $\widehat{\boldsymbol{S}} = \frac{\boldsymbol{S}}{l}$ and $\widehat{\boldsymbol{S}}^{\perp} = \hat{\mathbf{z}} \times \widehat{\boldsymbol{S}}$, which define the parallel and perpendicular axes of the spindle, respectively. The azimuthal angle in this frame of reference is given by $\theta_r$ (Fig S5A). In these coordinates, Eq. (10) is transformed into

$$\frac{\partial P^k}{\partial t} - \omega \frac{\partial P^k}{\partial \theta_r} = \Omega^k(\boldsymbol{y}, \boldsymbol{x}_k, \dot{\boldsymbol{x}}_k) \frac{1 - P^1 - P^2}{1 + (N-1)(1 - P^1 - P^2)} - \kappa P^k \tag{21}$$

Note that taking the limit $t \to \infty$ in this frame of reference is possible, since now $\boldsymbol{x}_k$ is not time dependent.

We change variables $P_s = P^1 + P^2$ and $P_d = P^1 - P^2$ , with redefined impingement rates $\Omega_s = \Omega^1 + \Omega^2$ and $\Omega_d = \Omega^1 - \Omega^2$. Also, to simplify the expression we define an interaction function

$$I(P_s) = \frac{1 - P_s}{1 + (N-1)(1 - P_s)} \tag{22}$$

We first take the limit of $t \to \infty$, then take derivatives with respect to $\omega$, and finally evaluate at $\omega = 0$, resulting in four equations for the unknowns $\bar{P}_s = P_s|_{\omega=0}$, $\bar{P}_{d,} = P_d|_{\omega=0}$, $\partial_\omega \bar{P}_s = \left.\frac{\partial P_s}{\partial \omega}\right|_{\omega=0}$ and $\partial_\omega \bar{P}_d = \left.\frac{\partial P_d}{\partial \omega}\right|_{\omega=0}$:

$$0 = \Omega_s(\boldsymbol{y}, \boldsymbol{x}_k, \dot{\boldsymbol{x}}_k) I(\bar{P}_s) - \kappa \bar{P}_s \tag{23}$$

$$0 = \Omega_d(\boldsymbol{y}, \boldsymbol{x}_k, \dot{\boldsymbol{x}}_k) I(\bar{P}_s) - \kappa \bar{P}_d \tag{24}$$

$$-\frac{\partial \bar{P}_s}{\partial \theta_r} = \left.\frac{\partial \Omega_s(\boldsymbol{y}, \boldsymbol{x}_k, \dot{\boldsymbol{x}}_k)}{\partial \omega}\right|_{\omega=0} I(\bar{P}_s) + \partial_\omega \bar{P}_s (I'(\bar{P}_s) \Omega_s(\boldsymbol{y}, \boldsymbol{x}_k, \dot{\boldsymbol{x}}_k)|_{\omega=0} - \kappa) \tag{25}$$

$$-\frac{\partial \bar{P}_d}{\partial \theta_r} = \left.\frac{\partial \Omega_d(\boldsymbol{y}, \boldsymbol{x}_k, \dot{\boldsymbol{x}}_k)}{\partial \omega}\right|_{\omega=0} I(\bar{P}_s) + \partial_\omega \bar{P}_d I'(\bar{P}_s) \Omega_d(\boldsymbol{y}, \boldsymbol{x}_k, \dot{\boldsymbol{x}}_k)|_{\omega=0} - \kappa \frac{\partial \bar{P}_d}{\partial \omega} \tag{26}$$

Note that $P_s, P_d, \frac{\partial P_s}{\partial \omega}$, and $\frac{\partial P_d}{\partial \omega}$ can be solved analytically, which results in a closed form for rotational drag coefficient

$$H_{CFG} = -M f_0 \frac{l}{4\pi R^2} \frac{1}{2} \int_{|y|=R} \frac{\widehat{\boldsymbol{S}}^\perp \cdot \boldsymbol{y}}{|\boldsymbol{y} - \boldsymbol{S}|} (\partial_\omega \bar{P}_s + \partial_\omega \bar{P}_d) - \frac{\widehat{\boldsymbol{S}}^\perp \cdot \boldsymbol{y}}{|\boldsymbol{y} + \boldsymbol{S}|} (\partial_\omega \bar{P}_s - \partial_\omega \bar{P}_d) dA(\boldsymbol{y}) \tag{27}$$

Note that this calculation is valid in shapes with axial symmetry. Since all the terms in the integrand are known, $H_{CFG}$ can be numerically approximated by standard quadrature techniques.

***Effective rotational drag: An asymptotic expansion for short spindles***

It is interesting to compute an approximation of $H_{CFG}$ when the spindle size is much smaller than the cell size. We define our perturbation parameter as $\delta := \frac{l}{R}$, and we determine $H_{CFG}$ up to the first order. We start by linearizing Eq. (26)

$$H_{CFG} = -Mf_0 \frac{l}{4\pi R^2} \int_{|y|=R} \widehat{\boldsymbol{S}}^{\perp} \cdot \widehat{\boldsymbol{y}} \left(\partial_\omega \bar{P}_d + \delta \widehat{\boldsymbol{S}} \cdot \widehat{\boldsymbol{y}}\ \partial_\omega \bar{P}_s + \mathcal{O}(\delta^2)\right) dA(\boldsymbol{y}) \tag{28}$$

Note that $\partial_\omega \bar{P}_s = \mathcal{O}(\delta)$, therefore the second term in the integral in Eq. (27) is $\mathcal{O}(\delta^2)$ and can be neglected. We do a non-standard multipole-type expansion of the impingement rate, using the classical expansions for $\frac{|x|}{|y|} < 1$:

$$\frac{e^{-\frac{|x-y|}{L}}}{|\boldsymbol{x}-\boldsymbol{y}|} = \frac{1}{L} \sum_{n\geq 0} (2n+1) i_n\left(\frac{|\boldsymbol{x}|}{L}\right) k_n\left(\frac{|\boldsymbol{y}|}{L}\right) P_n(\widehat{\boldsymbol{x}} \cdot \widehat{\boldsymbol{y}}) \tag{29}$$

$$\frac{\boldsymbol{y}-\boldsymbol{x}}{|\boldsymbol{x}-\boldsymbol{y}|^2} = \sum_{n\geq 0} \left(\frac{|\boldsymbol{x}|}{|\boldsymbol{y}|}\right)^n T_n(\widehat{\boldsymbol{x}} \cdot \widehat{\boldsymbol{y}}) \frac{\widehat{\boldsymbol{y}}}{|\boldsymbol{y}|} - \sum_{n\geq 0} \left(\frac{|\boldsymbol{x}|}{|\boldsymbol{y}|}\right)^{n+1} U_n(\widehat{\boldsymbol{x}} \cdot \widehat{\boldsymbol{y}})(I - \widehat{\boldsymbol{y}}\widehat{\boldsymbol{y}}) \frac{\widehat{\boldsymbol{x}}}{|\boldsymbol{y}|} \tag{30}$$

$$\frac{1}{|\boldsymbol{x}-\boldsymbol{y}|} = \frac{1}{|\boldsymbol{y}|} \sum_{n\geq 0} \left(\frac{|\boldsymbol{x}|}{|\boldsymbol{y}|}\right)^n P_n(\widehat{\boldsymbol{x}} \cdot \widehat{\boldsymbol{y}}) \tag{31}$$

where

- $i_n$ and $k_n$ are the modified Bessel function of the first and second kind.
- $P_n$ are the Legendre polynomials of the first kind.
- $T_n$ and $U_n$ are the Chebyshev Polynomials of the first and second kind.

Keeping the terms up to first order in $\delta$, we have the approximation

$$\Omega_s \approx \gamma e^{-\frac{R}{L}}\chi \tag{32}$$

$$\Omega_d \approx \gamma e^{-R/L}\chi\delta\left(\left(1 \, + \, 3\frac{i_1\left(\frac{l}{L}\right)k_1\left(\frac{R}{L}\right)\frac{R}{L}}{i_0\left(\frac{l}{L}\right)k_0\left(\frac{R}{L}\right)}\right)\widehat{\boldsymbol{\rho}} + \omega\frac{R}{V_g}\widehat{\boldsymbol{\theta}}\right)\cdot\widehat{\boldsymbol{y}} \tag{33}$$

However, in the limit when $l \, << \, L$, we can further approximate Eq. (32) by

$$\Omega_d \approx \gamma e^{-R/L}\chi\delta\left(\left(2 \, + \, \frac{R}{L}\right)\widehat{\boldsymbol{\rho}} + \omega\frac{R}{V_g}\widehat{\boldsymbol{\theta}}\right)\cdot\widehat{\boldsymbol{y}} \tag{34}$$

And so, up to first order we have

$$\frac{\partial P_d}{\partial\omega} \approx -\,\delta\,\left(\widehat{\boldsymbol{\theta}}\cdot\widehat{\boldsymbol{y}}\right)P_s^{(0)}\left(2 + \frac{R}{V_g}(\lambda - \kappa)\right)\frac{1}{\kappa} \tag{35}$$

where $P_s^{(0)}$ is the solution to Eq. (22) to 0-th order, given by

$$P_s^{(0)} = \frac{\frac{\gamma\chi e^{-\frac{R}{L}}}{\kappa} + N - \sqrt{\left(\frac{\gamma\chi e^{-\frac{R}{L}}}{\kappa} - N\right)^2 + 4\frac{\gamma\chi e^{-\frac{R}{L}}}{\kappa}}}{2(N-1)} \tag{36}$$

Note that, in the limit of $N \to 1$, $P_s^{(0)} \to \frac{\Omega}{\Omega+\kappa}$, while for $N \to \infty$, $P_s^{(0)} \approx \frac{\Omega}{\kappa N}$.

Finally, by defining the new variables:

1. $\tau_0 = lf_0$, which corresponds to the maximum torque from one motor.

2. $\bar{M} = P_s^{(0)}M$, which corresponds to the effective number of motors.

3. $\sigma = \frac{l}{3R}\left(2 + \frac{R}{V_g}(\lambda - \kappa)\right)$, which is a geometric factor relating microtubule dynamics and cell size.

By Eq. (27), the rotational drag coefficient of the spindle coming from microtubules is approximately

$$H_{CFG} \approx \tau_0 \bar{M} \sigma \frac{1}{\kappa} \tag{37}$$

The numerical approximation of Eq. (27) matches the first order analytical solution well (Figure S4A). Setting the half-spindle length of 5.5 µm, we find that $H_{CFG} = 8.03$ pN min µm / degree in the analytical and 13.30 pN min µm / degree in the numerical solutions.

***Simulations.***

In the main body of the text, we provide simulations of Eqs. (10, 12-15) (Fig 5, S5). All the effort lies in evaluating the integral in Eq. (12). To do this we discretize the surface of the sphere using the Lebedev quadrature with 434 points, with nodes and weights $\{\boldsymbol{y_i}, \boldsymbol{w_i}\}_{i=1}^{434}$; this quadrature exactly integrates the first 35 spherical harmonics [5]. Thus, we have a system of ODEs for the variables $\boldsymbol{x_C}, \boldsymbol{S}$, and $\{P^1(\boldsymbol{y_i}, \ \cdot\ ), P^2(\boldsymbol{y_i}, \ \cdot\ )\}_1^{434}$. We used 4th order Runge Kutta for evolving the system.

The quadrature for Eq. (27) is the same as that for Eq. (12). We performed a self-consistent convergence analysis of our quadrature method using Eq. (27), with the parameters used in this study. Using a very fine mesh of 5294 quadrature points as the exact solution, we find 9 digits of agreement.

**Table S1**. Model Parameters.

Note: $M, N$ and $r$ differed between our analyses of spindle movement (Fig 5F) and spindle rotational drag coefficients (Fig 5D,G,H,I, 6B, S4).

| **Parameter** | **Description** | **In Simulation** | **Source** |
|---|---|---|---|
| $R$ | Cell radius | 12 µm | Measured in this study |
| $2l$ | Spindle length (centrosome- | 11 µm | Measured in this study |

| | | | |
|---|---|---|---|
| | centrosome) | | |
| $\boldsymbol{x}_k$ | Position of *k*th centrosome | - | - |
| $\gamma$ | Microtubule nucleation rate | 1000 1/s | Based on [6] |
| $V_g$ | Microtubule growing speed | 2 μm/s | Measured in this study |
| $\lambda$ | Microtubule catastrophe rate | $\frac{V_g}{L}$ | - |
| $L$ | Average microtubule length | 5 μm | Estimate |
| $M$ | Total number of force generators | Varies | - |
| $r$ | Force generator capture radius | Varies | Estimated as the extent of microtubule length in contact with the cortex from EB1 imaging |
| $N$ | Number of force generators per site | Varies | - |
| $\frac{M}{N}$ | Force generator clusters | - | - |
| $f_0$ | Force generator pulling force | 1 pN | Based on [7-9] and measured in this study. |
| $\kappa$ | Rate of microtubule detachment from force generator | 1/15 1/s | Estimated based on EB1 comet persistence at cortex |
| $\mu_{cyto}$ | Translational drag coefficient due to fluid interactions of cytoplasm | 400 pN / s | Estimated with measurements in this study |
| $H_{cyto}$ | Rotational drag coefficient due to fluid interactions of cytoplasm | 1 pN μm min/degree | Estimated with measurements in this study |

**Supplementary Figure 1. Details on magnetic tweezer system.** (A) Schematic of the magnetic tweezer system, sample, and imaging setup. (B) Calibration mean-squared displacement (MSD) measurements of magnetic beads diffusing in pure glycerol (black boxes) with fitted line (solid blue). Dashed lines showing expected MSD in decreasing percentages of glycerol with water. (C) Sample 2D force calibration map around the tweezer tip (black triangle) showing forces from 0-35 pN. (D) Corresponding error on force map shown in C. (E) Example calibration trajectories of beads pulled through glycerol showing the difference between actual force used in the calibration measurement and the interpolated 2D force map. Most trajectories are well-fit by the calibration map (green, less than 20% off) with some outliers likely due to image segmentation artefacts (red, greater than 80% off).

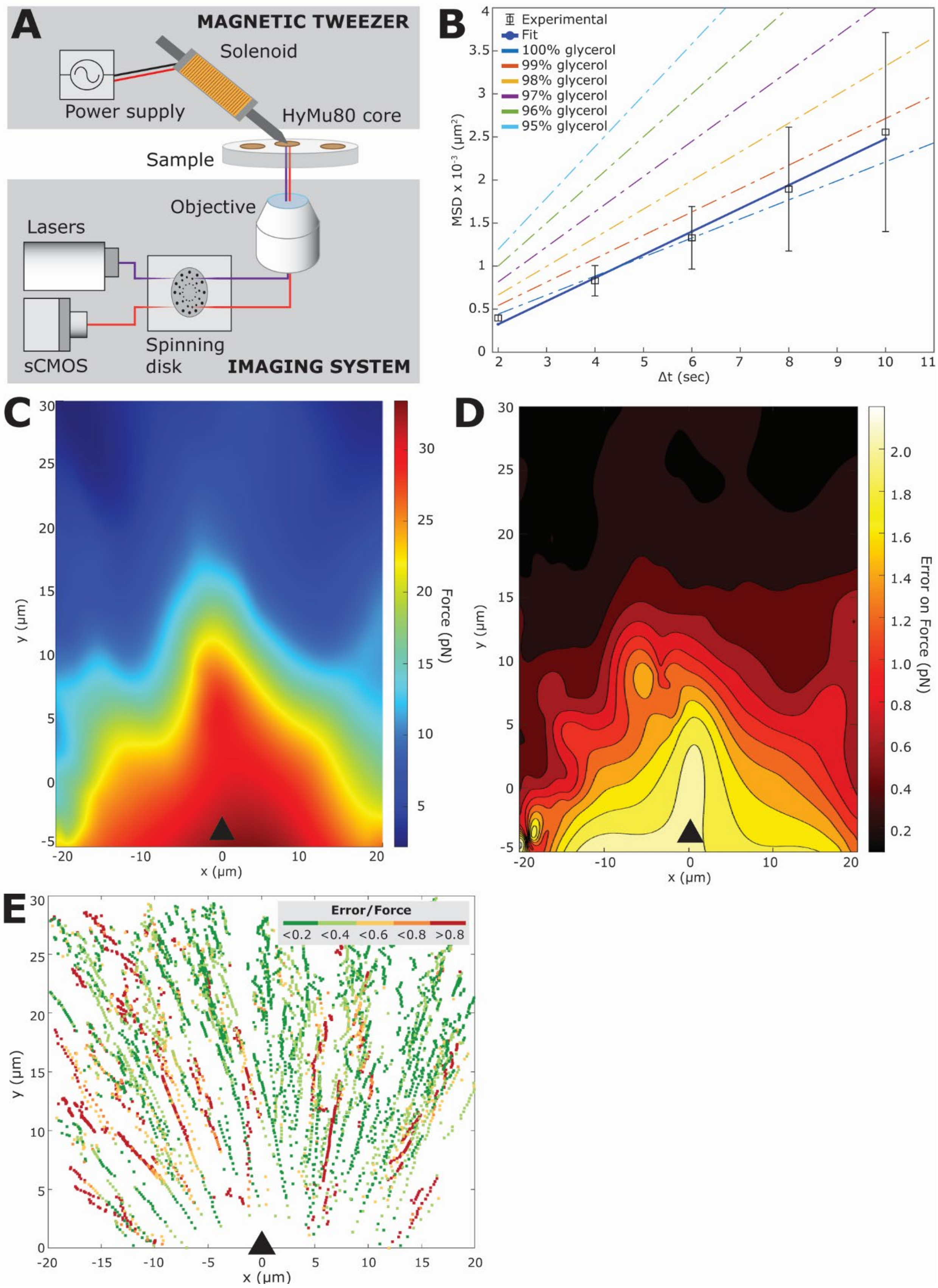
A
MAGNETIC TWEEZER
Solenoid
Power supply
HyMu80 core
Sample
Objective
Lasers
sCMOS
Spinning disk
IMAGING SYSTEM
B
Experimental
Fit
100% glycerol
99% glycerol
98% glycerol
97% glycerol
96% glycerol
95% glycerol
MSD x 10⁻³ (µm²)
Δt (sec)
C
Force (pN)
y (µm)
x (µm)
D
Error on Force (pN)
y (µm)
x (µm)
E
Error/Force
<0.2 <0.4 <0.6 <0.8 >0.8
y (µm)
x (µm)

**Supplementary Figure 2. Superparamagnetic beads can be introduced to cells without negatively impacting mitosis and daughter cell survival.** (A) Schematic of the procedure to introduce beads to cells. (B) Comparison of anaphase pole-pole separation shows that the presence of beads does not significantly impact cell division (compare control, brown, n=13 to single beads, light brown, n=7 or multiple beads, light blue, n=8). Anaphase pole separation is slightly slower after pushing on the spindle (single beads after push, dark green, n=5), although cells still divide successfully. (C) Time course of U2OS cell labeled with GFP-hCentrin2, GFP-CENPA, and mCherry-alpha Tubulin in early anaphase showing that the bead (cyan, 647-N biotin) is easily moved around by the spindle during division. Scale bar is 5 μm. (D) Time course of late anaphase shows a cleavage furrow beginning to physically divide daughter cells with one daughter containing the bead. Scale bar is 5 μm.

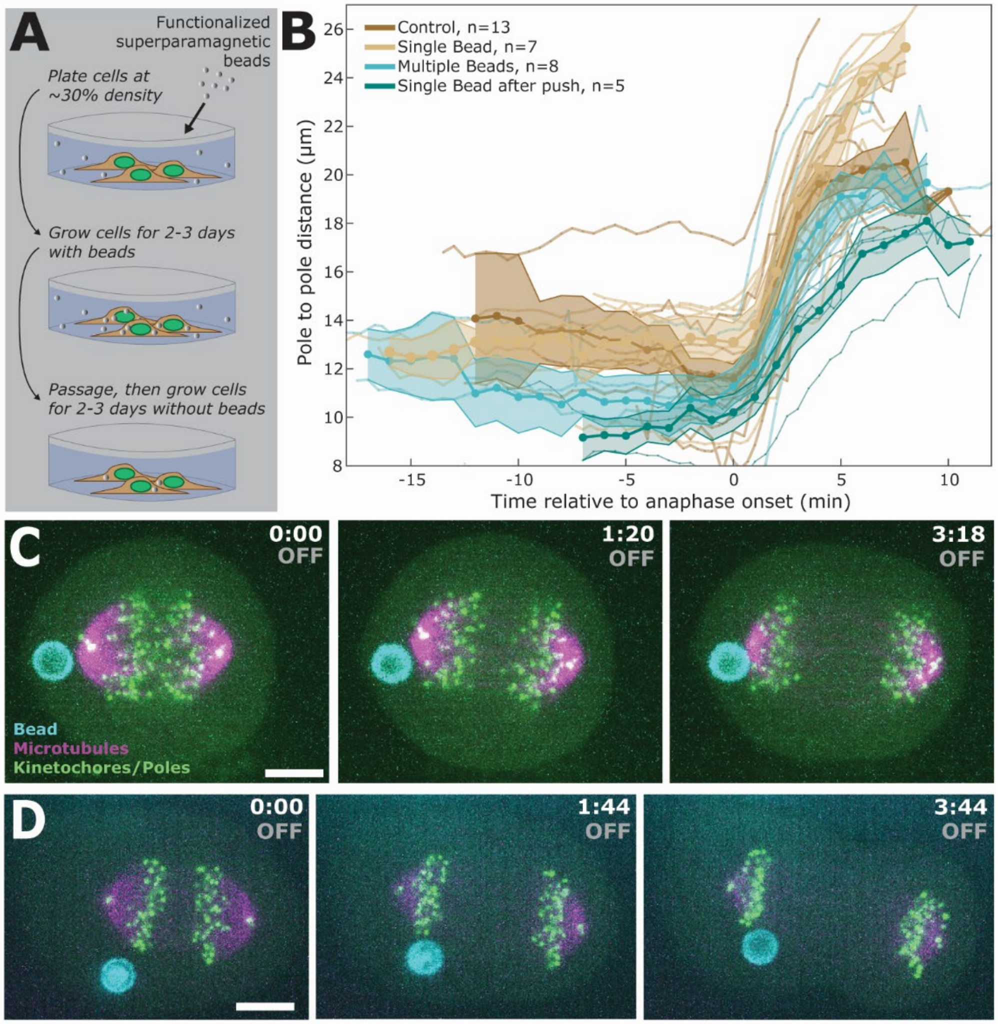
A
Functionalized superparamagnetic beads
Plate cells at ~30% density
Grow cells for 2-3 days with beads
Passage, then grow cells for 2-3 days without beads
B
Control, n=13
Single Bead, n=7
Multiple Beads, n=8
Single Bead after push, n=5
Pole to pole distance (μm)
Time relative to anaphase onset (min)
C
0:00 OFF
1:20 OFF
3:18 OFF
Bead
Microtubules
Kinetochores/Poles
D
0:00 OFF
1:44 OFF
3:44 OFF

**Supplementary Figure 3. Additional phenotypic characterization.** (A) RNAi of LGN (green, n=22) does not affect metaphase spindle length, anaphase pole separation velocity, or final anaphase spindle length compared to WT (grey, n=17). (B) RNAi of individual gαs (gα1/3, n=22 or gα2, n=26) or all gαs (gα1/3 + gα2, n=28) do not affect metaphase spindle length, anaphase pole separation velocity, or final anaphase spindle length compared to WT. (C) RNAi of individual gαs does not change cortical dynein localization as shown with representative images of U2OS cells expressing tdTomato-DYNHC2 (pink) and GFP-hCentrin2 (green). (D) Neither RNAi of gα1/3 nor of gα2 alone significantly changed the spindle's rotational drag (n=10, p=0.94 and n=10, p=0.86, respectively). Further, both RNAi of gα1/3 and gα2 alone had significantly higher rotational drag than the combined gα RNAi treatment (p=0.04 and 0.04, respectively) (E) Example timepoints of cortical EB1 comets at the coverslip plane are shown (white arrows showing 2 microtubule tips that move between the top and bottom frames). (F) RNAi of DLG1 does not significantly change metaphase spindle length, anaphase pole separation velocity, or final anaphase spindle length (n=22). RNAi of MARK2 decreases pole separation velocity (n=20). (G) RNAi of KIF13B does not impact the spindle's rotational drag (n=8, p=0.16). A double RNAi treatment of LGN and KIF13B (n=9) shows a similar decrease from WT rotational drag as LGN alone (p=0.003 compared to WT). (H) Treatment with both DLG1 and MARK2 siRNAs resulted in a decreased rotational drag coefficient (1.7 ± 0.5 pN min µm / degree, n=25) compared to MARK2 (p=0.004) and WT (p=0.006), but no difference from DLG1 alone (p=0.06).

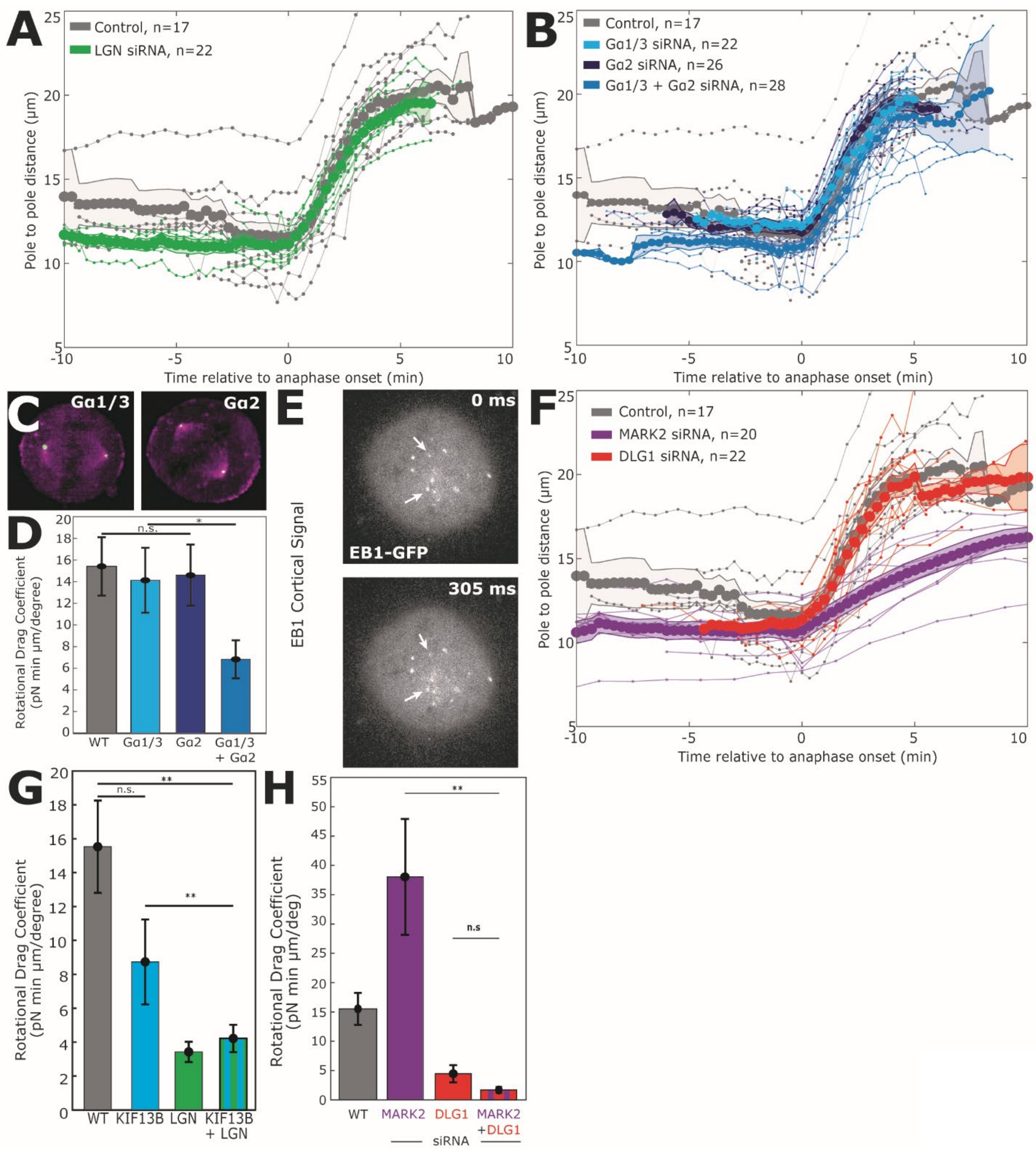

A
Control, n=17
LGN siRNA, n=22
Pole to pole distance (μm)
Time relative to anaphase onset (min)
B
Control, n=17
Gα1/3 siRNA, n=22
Gα2 siRNA, n=26
Gα1/3 + Gα2 siRNA, n=28
Pole to pole distance (μm)
Time relative to anaphase onset (min)
C
Gα1/3
Gα2
D
Rotational Drag Coefficient (pN min μm/degree)
n.s.
*
WT
Gα1/3
Gα2
Gα1/3 + Gα2
E
EB1 Cortical Signal
0 ms
EB1-GFP
305 ms
F
Control, n=17
MARK2 siRNA, n=20
DLG1 siRNA, n=22
Pole to pole distance (μm)
Time relative to anaphase onset (min)
G
Rotational Drag Coefficient (pN min μm/degree)
n.s.
**
**
WT
KIF13B
LGN
KIF13B + LGN
H
Rotational Drag Coefficient (pN min μm/deg)
**
n.s
WT
MARK2
DLG1
MARK2 +DLG1
siRNA

**Supplementary Figure 4. Dynein and LGN form clusters at the cortex.** (A) (top) WT dynein cortical signal is patchy within the midline crescents. Scale bar is 10 µm. (bottom) Higher magnification shows evidence of clustering within the midline crescent. Scale bar is 5 µm. (B) (top) WT LGN signal is patchy within midline crescents. Scale bar is 10 µm. (middle) Higher magnification shows evidence of clustering, similar to the dynein phenotype. Scale bar is 5 µm. (bottom) LGN forms spots visible at the coverslip, suggesting that patchiness in the midline crescents is due to imaging slices of cortical clusters. Scale bar is 5 µm. (C) Schematic of dynein and LGN clustering at the cortex.

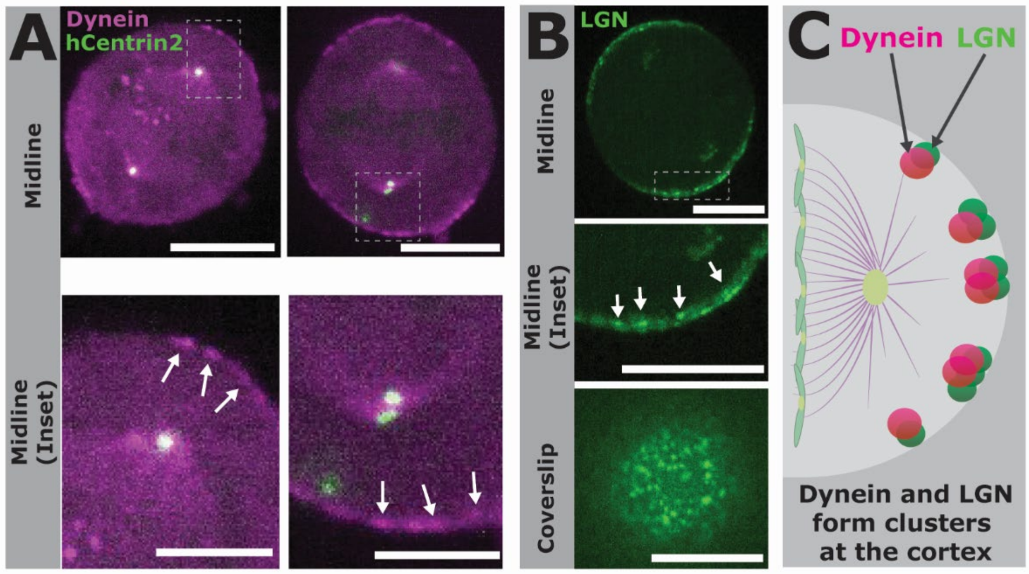

**Supplementary Figure 5. Effect of varying model parameters on the spindle's rotational drag coefficient.** (A) Schematic of rotating reference frame centered on the spindle axis, showing vectors $\widehat{\boldsymbol{S}} = \frac{\boldsymbol{s}}{l}$ and $\widehat{\boldsymbol{S}}^{\perp} = \hat{\boldsymbol{z}} \times \widehat{\boldsymbol{S}}$ which define the parallel and perpendicular axes of the spindle, respectively. The azimuthal angle in this frame of reference is given by $\theta_r$. (B) Comparison of numerical and first order analytical solutions of the torque exerted on the spindle by bound microtubules. (C) Increasing astral microtubule length results in an increased rotational drag coefficient. (D) Increasing force generator radius results in increased rotational drag on the spindle. (E) Decreasing the microtubule catastrophe rate results in higher rotational drag.

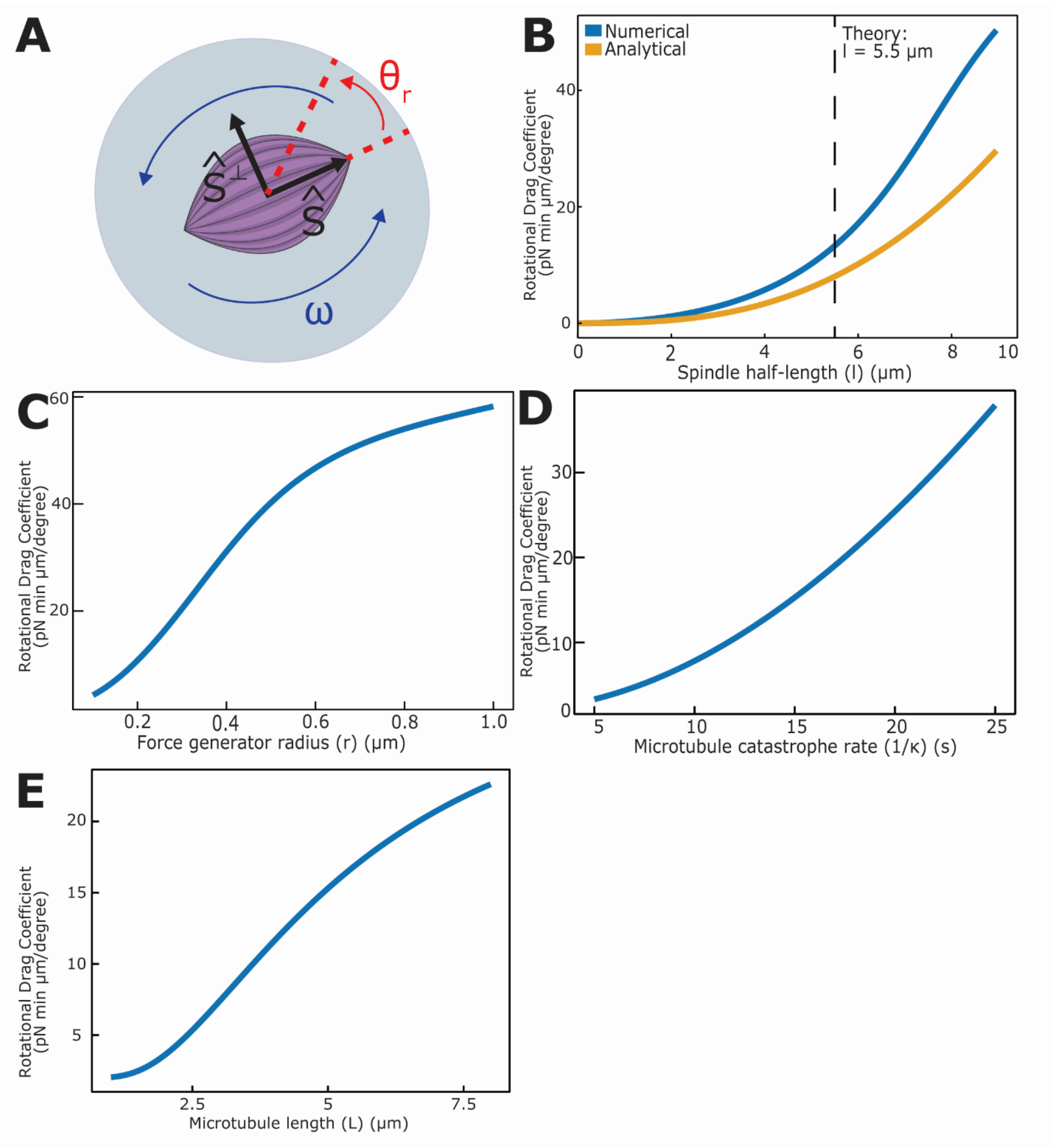
A
θr
Ŝ⊥
Ŝ
ω
B
Numerical
Analytical
Theory:
l = 5.5 μm
Rotational Drag Coefficient (pN min μm/degree)
Spindle half-length (l) (μm)
C
Rotational Drag Coefficient (pN min μm/degree)
Force generator radius (r) (μm)
D
Rotational Drag Coefficient (pN min μm/degree)
Microtubule catastrophe rate (1/κ) (s)
E
Rotational Drag Coefficient (pN min μm/degree)
Microtubule length (L) (μm)
